# Maternal body condition and season influence RNA deposition in the oocytes of alfalfa leafcutting bees (*Megachile rotundata*)

**DOI:** 10.1101/2022.10.11.511817

**Authors:** Mallory A. Hagadorn, Frances K. Hunter, Tim DeLory, Makenna M. Johnson, Theresa L. Pitts-Singer, Karen M. Kapheim

**Author notes:** These authors contributed equally to this work and share first authorship (Mallory A. Hagadorn, Frances K. Hunter).

## Abstract

Maternal effects are an important source of phenotypic variance, whereby females influence offspring developmental trajectory beyond direct genetic contributions, often in response to changing environmental conditions. However, relatively little is known about the mechanisms by which maternal experience is translated into molecular signals that shape offspring development. One such signal may be maternal RNA transcripts (mRNAs and miRNAs) deposited into maturing oocytes. These regulate the earliest stages of development of all animals, but are understudied in most insects. Here we investigated the effects of female internal (body condition) and external (time of season) environmental conditions on maternal RNA in the maturing oocytes and 24 hr old eggs of alfalfa leafcutting bees. Using gene expression and WGCNA analysis, we found that females adjust the quantity of mRNAs related to protein phosphorylation, transcriptional regulation, and nuclease activity deposited into maturing oocytes in response to both poor body condition and shorter day lengths that accompany the late season. However, the magnitude of these changes was higher for time of season. Females also adjusted miRNA deposition in response to seasonal changes, but not body condition. We did not observe significant changes in maternal RNAs in response to either body condition or time of season in 24-hr-old eggs, which were past the maternal-to-zygotic transition. Our results suggest that females adjust the RNA transcripts they provide for offspring to regulate development in response to both internal and external environmental cues. Variation in maternal RNAs may, therefore, be important for regulating offspring phenotype in response to environmental change.

## Introduction

Females can influence offspring development in ways that are independent of direct genetic inheritance, typically in response to changing environmental cues (Bernardo 1996*a*, Wolf & Wade 2009). The mechanisms underpinning such maternal effects include both pre- and post-zygotic functions. For example, females can influence the developmental rate, size, or sex of their offspring through post-zygotic mechanisms such as choice of nest site, incubation frequency, or provisioning rate (Meaney 2001, Bernardo 1996*a*, Torchio & Tepedino 1980, Klostermeyer et al. 1973). Females can also influence the earliest stages of offspring development by adjusting the transcriptional and endocrine profiles of maturing oocytes (Groothuis et al. 2019, Vastenhouw et al. 2019, Wolf & Wade 2009). Although these pre-zygotic mechanisms are likely to have early and ongoing effects on offspring phenotype, they have been relatively understudied compared to post-zygotic mechanisms, especially in insects. This is particularly true with regard to how females interpret environmental cues and translate them into molecular signals that influence offspring development (Lee & Duvall 2022, Huestis & Marshall 2006). Furthermore, although it is likely that maternal effects are influenced by multiple cues (Marshall & Uller 2007), these cues are typically studied in isolation (though see Potticary & Duckworth (2020)). Studies assessing the effects of multiple cues—a more realistic view of environmental influences—will yield a better understanding of non-genetic drivers of phenotypic variance, which provide the raw material for adaptive evolution and are an important source of ecological diversity (Mousseau & Dingle 1991, Mousseau & Fox 1998, Bernardo 1996*a*, Räsänen & Kruuk 2007).

Cues influencing pre-zygotic maternal effects on offspring phenotype can come from the external or internal environment. External cues experienced by most females are both biotic and abiotic, including interactions with natural enemies (Agrawal et al. 1999, Sharda et al. 2021, Mitchell & Read 2005, Rolff 1999) and exposure to changing weather conditions (Burgess & Marshall 2011, Bernardo 1996*a*). Photoperiod is a common environmental cue, because it can serve as a reliable indicator of seasonal change (Bradshaw & Holzapfel 2007), and can thus serve as a coordinating mechanism for critical life history events such as reproduction, mating, migration, and diapause (Bradshaw & Holzapfel 2007, Mousseau & Dingle 1991). For example, maternal photoperiod influences egg size, diapause, and survival outcomes of offspring in mosquitoes (Lacour et al. 2014, Lee & Duvall 2022). Similarly, maternal photoperiod is the primary determinant of whether larvae enter diapause in multiple fly species (Saunders et al. 1986, McWatters & Saunders 1997). There is also evidence for transgenerational effects of photoperiod on offspring diapause, development time, and several morphological traits in a parasitoid wasp (Tougeron et al. 2020). Together, the results of these studies suggest female response to external cues such as photoperiod is an important driver of maternal effects in insects.

Environmental cues that shape maternal effects may also stem from changes in one or more internal conditions. These may include hormones (for reviews see Edwards et al. 2021, Groothuis & Schwabl 2008, Groothuis et al. 2019, Meylan et al. 2012), body size (Steiger 2013), and body condition (de Zwaan et al. 2019). Interestingly, maternal nutritional status can also impact offspring development. In non-biting midges, females reared in food stressed conditions yielded offspring that developed faster and had decreased fecundity relative to those given resources in excess (Colombo et al. 2014). Similar effects of parental nutritional status have been observed in mosquitoes, where offspring of nutritionally-stressed parents were more likely to transmit the dengue virus (Zirbel et al. 2018). In neriid flies, offspring of females in poor body condition developed faster than those from females in good body condition (Bonduriansky & Head 2007). Finally, parasitoid wasps experiencing high levels of competition, which may decrease body condition, were more likely to produce diapausing offspring (Tougeron et al. 2018). Based on these studies, it is clear that females can influence the same set of offspring traits (e.g., development rate, fecundity, and even diapause) in response to different sets of internal and external environmental cues. How females integrate these cues to influence offspring development via maternal effects is unknown. Resolving this relationship will require perspective on how cues from the environment are translated into signals that regulate development in offspring.

Understanding how maternal experience (i.e., internal and external cues) is translated into variation in offspring phenotype requires insight into the mechanisms by which females can influence the developmental trajectory of their offspring. One of the earliest stages at which females can influence offspring phenotype is during oogenesis. During this stage, small shifts in maternal input of hormones (Navara et al. 2006), yolk (or nutrition) (Bernardo 1996*b*), and RNA (Vastenhouw et al. 2019) can have large, organizational effects on the developing zygote. In insects, maternal RNAs are transcribed in the supporting nurse cell and transferred into the growing oocyte via cytoskeletal machinery (Spradling 1993). Maternal RNAs can include both protein-coding (mRNA) and small, non-coding regulatory RNAs (e.g., microRNA [miRNA]), and are necessary for the final stages of oocyte maturation and activation and the earliest stages of embryonic development following fertilization (Winata & Korzh 2018, Vastenhouw et al. 2019). These maternally-derived molecules are responsible for regulating critical processes in early embryogenesis such as cell structure and division, biosynthesis, blastula formation, and gastrulation (Weeks & Melton 1987, Winata & Korzh 2018, Tang et al. 2007, Tadros & Lipshitz 2009, Pauli et al. 2011, Paranjpe et al. 2013, Torres-Paz et al. 2019, Baroux et al. 2008, Harvey et al. 2013). Maternal RNAs are programmatically degraded and cleared in a dynamic process leading up to the maternal-to-zygotic transition (MZT), which occurs during the blastoderm stage in insects (Pires et al. 2016, Sung et al. 2013). After the MZT, developmental processes, including RNA transcription, are controlled by the embryo.

Given their influential role in the earliest stages of development, variation in maternal RNAs are a potential way that females translate environmental cues into maternal effects on offspring phenotype. Research with fish has revealed extensive variation in how females deposit RNA transcripts in their eggs. For example, there are significant differences in the relative abundance of maternally-deposited transcripts among zebrafish females, but almost no variation within a female’s clutch (Rauwerda et al. 2016). This could suggest that female zebrafish make consistent adjustments in how they deposit transcripts during oogenesis based on their internal condition. In support of this, bacterial supplementation treatments in zebrafish modified maternal condition and influenced the composition of maternal mRNAs (Miccoli et al. 2017, 2015). Additionally, in the round goby, females adjust their RNA contributions to embryos in response to water temperature (Adrian-Kalchhauser et al. 2018). In the annual killifish, females respond to environmental cues via maternal programming of mRNAs and miRNAs deposited in oocytes that determine whether young embryos develop directly or enter diapause (Romney & Podrabsky 2017). Lastly, in both cichlids and cavefish, differences in maternal RNA provisioning play a causal role in generating the phenotypic novelty that promotes local adaptation and species diversification (Ahi et al. 2018, Torres-Paz et al. 2019). Overall, the potential for fish to alter offspring development and impact phenotype through environmentally-sensitive maternal RNA provisioning is evident.

There is accumulating evidence that insects also transmit signals about the environment to developing offspring via maternally-derived RNAs. Egg diapause in the Asian tiger mosquito (*Aedes albopictus*) is maternally regulated based on exposure of adults to short day lengths (Mori et al. 1981, Wang 1966). Experimental studies have demonstrated that females adjust the composition of mRNAs (Poelchau et al. 2011), but not miRNAs (Batz et al. 2017) deposited into mature oocytes in response to photoperiod regime, and these mRNA adjustments coincide with an increased likelihood of diapause among offspring (Poelchau et al. 2011). Further research suggests these differences in maternally-deposited mRNAs may lead to increasingly divergent expression profiles between diapause-destined and non-diapaused destined embryos, even past the MZT (Poelchau et al. 2013). Similarly, mRNA differences have been found in the ovaries of female locusts (*Locusta migratoria*) exposed to short- and long-days (Hao et al. 2019), which is known to trigger maternally-mediated diapause in locust eggs (Tanaka 1994). This suggests that maternally provisioned RNAs may be a common factor in the initiation of diapause in the egg stage. However, insects can diapause during any stage of the life cycle and there is a diverse array of molecular signals that trigger diapause across species (Denlinger 2002, 2022). A complete understanding of maternal effects on insect diapause requires investigating species that diapause at different life stages.

We investigated the mechanistic underpinnings of maternal effects in alfalfa leafcutting bees (*Megachile rotundata*). Various aspects of alfalfa leafcutting bee biology make them an ideal species for investigating how maternal perception of the environment stemming from internal and external cues can shape maternal effects on offspring development. First, *M. rotundata* naturally exhibit facultative diapause as late-stage larvae (“prepupae”) (Pitts-Singer 2020, Pitts-Singer & Cane 2011, Tepedino & Parker 1988, 1986, Krunic 1972, Hobbs & Richards 1976); hence, a readily-observable dichotomy in developmental trajectory already exists among offspring. Second, maternal influence has long been recognized as a factor driving this facultative diapause (Tepedino & Parker 1986, Parker & Tepedino 1982, Johnson 2022). Therefore, their life cycle offers a unique opportunity for evaluating how maternal environmental cues are transmitted to the offspring. Third, two maternal cues, photoperiod and lipid stores (i.e., external and internal, respectively) are known to influence the probability of offspring diapause (Johnson 2022, Pitts-Singer 2020, Wilson et al. 2021). Adult *M. rotundata* females experience seasonal changes (early to late summer) in photoperiod, and manipulations of maternal day length experience has been shown to alter patterns of diapause induction in the offspring (Pitts-Singer 2020, Wilson et al. 2021). Likewise, maternal condition is also a significant determinant of diapause fate in progeny (Johnson 2022). Females regulate offspring size via the amount of food provided in the brood cell (Klostermeyer et al. 1973), which impacts the probability of survival during diapause (Fischman et al. 2017). Because brood cell construction and provisioning is energetically demanding (Klostermeyer & Gerber 1969, Klostermeyer et al. 1973), maternal body condition is likely to influence offspring diapause destiny. Jointly, these features make *M. rotundata* useful for disentangling how females integrate external and internal cues of the environment to direct offspring development via maternal RNA.

To test how external signals and internal condition influence the mechanisms of maternal effects, we explored the impacts of time of season (i.e., photoperiod) and maternal body condition (i.e., depletion of lipid stores) on maternal RNA in *M. rotundata*. Here, time of season includes the combined effects of all changes that occur during a season, including photoperiod, but also pests, temperature, floral resources, etc. To determine whether and how these cues influenced maternal RNAs, we quantified mRNA and miRNA from maturing oocytes and eggs that were 24 hours post-oviposition, and, therefore, past the maternal-to-zygotic transition (hereafter referred to as “24-hr eggs”). In our previous research, we found that females with experimentally reduced lipid stores had fewer diapausing offspring in both the early season and the late season. However, the probability of offspring diapause was significantly higher in the late season than in the early season. We thus hypothesized that *M. rotundata* females manipulate their RNA provisions as a response to both external and internal cues of the environment in a way that is consistent with known maternal effects on offspring diapause outcomes. Our results reveal a potential mechanism for observable maternal effects in bees.

## Methods

### Study Organism and Field Collections

*Megachile rotundata* are solitary bees that are intensively managed for alfalfa seed production (Pitts-Singer & Cane 2011, Calderone 2012, Pitts-Singer 2020). They readily nest in above-ground, artificial tunnels made of wood or polystyrene when they are used as commercial pollinators (Fairey & Lieverse 1986, Pitts-Singer & Cane 2011). In the western U.S., most offspring overwinter as cocooned, non-feeding fifth instar larvae (prepupae), but almost half the progeny avert this process, emerging as adults in the same summer as the parent generation (Pitts-Singer 2020, Pitts-Singer & Cane 2011, Tepedino & Parker 1988, 1986, Krunic 1972, Hobbs & Richards 1976).

For this study, we received bee cells containing diapausing prepupae from farms in the alfalfa seed-growing areas of Box Elder County, Utah, USA during October 2019 and kept them in cold storage at 4°C until diapause was broken by incubating the cocoons at 29°C. To manipulate female body condition, we interrupted development during incubation by temporarily moving bees into cool storage (18°C) for either 1 d (control) or 14 d (poor condition) before returning them to 29°C until adult emergence. This type of interruption is a commonly used management technique for aligning the timing of bee emergence and field release with alfalfa bloom (Richards 1984). Previous research has demonstrated that bees can survive a prolonged setback in temperature for two weeks at 18°C (Yocum et al. 2010). Our recent research demonstrated that this developmental pause significantly reduces lipid quantity at emergence, and females who have experienced this interruption produce significantly fewer diapausing offspring and provide smaller provision masses for offspring than females that experience only a one-day interruption (Johnson 2022). Additionally, late season females produce significantly more diapausing offspring than early season females, most of whose offspring are non-diapausers (Johnson 2022). To assess seasonal effects at the molecular level, we released bees from each body condition treatment in both the early and late summer. Average day length was 15.22 minutes longer during our early season experiment (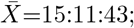 range=15:11:48–15:11:23) relative to the late season 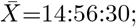 range=14:59:13–14:52:15).

Our experimental design allowed the release of females from both the poor body condition and control treatments in both early and late season. On the day of emergence, adult females were given a unique thoracic paint mark (Testors, Rockford, IL, USA) and then released into two 6.1 × 6.1 × 1.8 m screened cages erected over flowering alfalfa in Logan, Utah, USA. For both early and late season releases, all bees were placed into their respective cages within a 1-2 d period of each other. Male *M. rotundata* were released simultaneously in a ratio of 2:1 male to female (Rossi et al. 2010).

Each cage was provided with a small section of prefabricated, polystyrene nesting block mounted 1.1 m above the ground in the center of the cage. This allowed us to easily observe nesting activity. Cage assignments were random, but with equal numbers of control and poor body condition treatments in each cage. To compare how maternal RNA abundances respond to variation in body condition and time of season, we collected maturing oocytes and 24-hr eggs that were past the MZT. Nests were monitored three times daily to verify the identity of each nest owner. Additionally, we conducted hourly nest checks between 10:00 and 19:00 to see if any new eggs had been laid or were about to be laid. These hourly checks included watching for signs of nest initiation and pollen provisioning that are done just before egg laying. If an egg was laid, but the cell holding the egg not fully closed, we waited to collect the female until after she capped the cell. Once a female was known to have laid an egg, she was captured as soon as possible and flash frozen in liquid nitrogen within minutes. We later dissected the most mature oocyte from her ovaries (see below). Eggs still in the nest were left undisturbed for 24 hr, then frozen in liquid nitrogen, and later removed from the nest on dry ice prior to RNA isolation. We chose 24 hr for egg collection as this is post-MZT transition for insect eggs, including honey bees (Ninova et al. 2016, Tadros & Lipshitz 2009, Vastenhouw et al. 2019, Pires et al. 2016). These experimental methods provided us with maturing stage 4 oocytes and 24 hr eggs from females who experienced experimental lipid reduction and control females, in both the early and late season.

### Sample Processing

All samples were maintained at -80°C until dissection. Prior to dissections, abdomens were detached from the thorax while on dry ice and incubated in pre-chilled RNAlater-ICE Frozen Tissue Transition Solution (Thermo Fisher Scientific, Waltham, MA, USA) at -20°C. After a 16-20 hr incubation period, abdomens were dissected in RNAlater-ICE using a Leica M80 stereomicroscope with an IC80HD camera attached (Leica Microsystems, Buffalo Grove, IL, USA). While dissecting, we identified the longest terminal oocyte, removed it from the ovary, and transferred the oocyte to dry ice. Dissected oocytes were stored at -80°C until RNA isolation. Oocyte, trophocyte, and total egg chamber length was measured with the Leica Application Suite (v.4.5) software. We calculated the proportion of the egg chamber occupied by the oocyte and used these percentages, along with various egg chamber characteristics, to assign oocyte maturation stage as described by Kapheim & Johnson (2017). We selected oocytes from the same stage of maturation for each experimental group. This yielded 22 stage 4 oocytes occupying 79-95% of the egg chamber: ‘early control’, 85.8-92.8%, n=6; ‘early poor condition’, 78.9-89.7%, n=4; ‘late control’, 84.0-94.6%, n=6; ‘late poor condition’, 84.1-94.9%, n=6.

We isolated total RNA from a single oocyte (*n* = 22) and egg (*n* = 22) from each female in our study to yield 44 total samples. We used the *mir*Vana miRNA Isolation Kit with phenol (Ambion, Austin, TX, USA) according to the manufacturer’s protocol. The total RNA yield of isolates ranged from 0.52–1.54 *µ*g 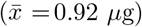 in oocytes to 0.54–1.27 *µ*g 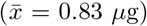 in eggs. Oocyte/egg pairs from the same female were always included in the same isolation batch, and we stored RNA isolates at -80°C until sequencing. RNA quality assessment was conducted by the Utah State University Center for Integrated Biosystems using a TapeStation 2200 with high sensitivity RNA reagents (Agilent Technologies, Santa Clara, CA, USA). Total RNA was sent to the Roy J. Carver Biotechnology Center at the University of Illinois at Urbana-Champaign for library preparation and sequencing.

All dissections, ovary measurements, and RNA isolations were conducted without knowledge of sample experimental treatment group.

### Sequencing

#### mRNA

RNAseq libraries were prepared with the Illumina TruSeq Stranded mRNAseq Sample Prep kit (Illumina). The libraries were pooled, quantitated by qPCR, and sequenced on two SP lanes for 151 cycles from both ends of the fragments on a NovaSeq 6000. This generated 1,832,084,298 total 100 nt reads with a mean of 41,638,279.50 (*±* 579,462.15 standard error) reads per sample. Fastq files were generated and demultiplexed with the bcl2fastq v2.20 Conversion software (Illumina).

#### miRNA

SmallRNA libraries were prepared using the Qiagen Small RNA Sample Prep kit. Libraries were pooled, quantitated by qPCR and sequenced on one SP lane for 51 cycles from one end of the fragments on a NovaSeq 6000. This generated 519,046,055 total 50 nt reads with a mean of 11,796,501.25 (*±* 248,977.88 standard error) reads per sample. Fastq files were generated and demultiplexed with the bcl2fastq v2.20 Conversion Software (Illumina). The 3’ adapters were trimmed.

### Alignment and Quantification

#### mRNA

After quality checks with MultiQC v1.5 (Ewels et al. 2016), we trimmed reads with Trimmomatic v.0.39 (Bolger et al. 2014) in Paired-End mode. We used the Illuminaclip function to remove adapters and the sliding window function to trim reads with an average quality score below 20, based on 4 nt windows. We used the ‘keep both reads’ option and filtered reads shorter than 51 nts. Trimmed reads were then aligned to the *M. rotundata* genome (Kapheim et al. 2015) using STAR v2.7.8a (Dobin et al. 2013) in paired end multi mode. We did not clip any bases from the read ends during alignment. Finally, we quantified the number of reads aligned to each CDS feature in the *M. rotundata* annotation v.1.1 (Kapheim et al. 2015) using the Python v3.6.3 script HTSeq-count v0.9.1 (Anders et al. 2015) in union mode with the stranded option set to reverse. The selection of these pipeline tools was made based on a comparative analysis of performance (Corchete et al. 2020).

#### miRNA

After quality checks with MultiQC v1.5 (Ewels et al. 2016), we implemented miRDeep2 for the quantification, alignment and identification of miRNA’s in *M. rotundata*. First, the *M. rotundata* genome (Kapheim et al. 2015) was indexed using bowtie-build. Then we ran each sample’s fastq miRNA through the *mapper*.*pl* script to align them to the *M. rotundata* genome. Reads shorter than 18 bases were discarded, non-nucleotide letters were removed (anything other than the following characters: ATCGUNatcgun), and reads were then collapsed. Next, we identified known miRNAs and discovered novel miRNAs using known precursor sequences from *M. rotundata*, and the known mature sequences from *M. rotundata, Bombus impatiens, B. terrestris, Apis mellifera, Megalopta genalis* and *Nomia melanderi* (Kapheim et al. 2020) as inputs alongside our sample sequences to the *miRDeep2*.*pl* script of miRDeep2 (Friedländer et al. 2011). Next, we ran the *quantifier*.*pl* script of miRDeep2 on found known and novel miRNAs for each *M. rotundata* individual sample to obtain read quantities for each pair of precursor and mature miRNA. We then filtered precursor-mature pairs to keep only those that were not rRNA/tRNAs, had a minimum of five reads each on mature and star strands of the hairpin sequence, and a randfold *p <* 0.05, following Kapheim et al. (2020). We then re-ran *miRDeep*.*pl* and *quantifier*.*pl* on the original sample set of mapped miRNAs. For the second run, we first compiled a list of the unique novel precursor miRNA sequences from across all samples which passed the aforementioned filtering criteria for the output from the first *quantifier*.*pl* run. This list of novel precursor miRNAs was added to the list of known precursor miRNAs of *M. rotundata* from Kapheim et al. (2020), and the composite was used as the species-specific precursor miRNA input for the second run of *miRDeep2*.*pl*, without changing any other inputs. We then applied the same filters to compile our final list of miRNAs.

For downstream analysis of differential miRNA expression, we treated miRNAs with identical mature sequences, but non-overlapping precursor locations, as separate miRNAs. Overlapping precursors with identical mature sequences were merged and considered the same miRNA. (Mapping to multiple precursor sequences did not occur within individuals.) miRNA homologs were identified with miRBase v22.1 (Griffiths-Jones et al. 2006). We used the search tool, powered by RNA Central, to identify the top hit for each mature miRNA sequence. We kept only those hits with query and target sequence match *>* 90% and no more than 2 mismatches, as long as those mismatches were not in the seed sequence region (2-7 nucleotides from the 5’ end) (Ambros et al. 2003, Griffiths-Jones et al. 2006).

### Target prediction of miRNAs

We ran the target prediction software, miRanda v3.3 (Enright et al. 2003), on the final set of unique mature miRNAs from all samples (minimum energy threshold -20, minimum score 140, strict alignment to seed region -en -20 -sc 140 -strict). In addition, we analyzed this same set of mature miRNAs for potential target sequences using RNAhybrid v2.12 (Krüger & Rehmsmeier 2006) (minimum free energy threshold -20). We kept only miRNA-target gene pairs that were predicted by both programs with *p <* 0.01.

### Statistical Analysis

#### mRNA

We combined all feature counts from the HTSeq-count output in R v4.0.2 (R Core Team 2020) and used these to make a DGEList with the EdgeR package v3.30.3 (Robinson et al. 2010). Visual inspection of samples on a PCA (Fig. S1) revealed clear separation between oocyte and egg samples and the presence of one outlier (egg sample 202198). We therefore proceeded to analyze these separately after removing the outlier. For each dataset (oocyte and egg), we used the *filterByExpr* function to filter genes based on counts per million (CPM) and number of samples in each group. This kept 8,276 (64.8%) and 9,200 (72.0%) out of 12,770 genes for oocytes and eggs, respectively. This concurs with the proportion of protein-coding genes known to be maternally transcribed and deposited into maturing oocytes in *Drosophila* (Laver et al. 2015). Libraries were then normalized with the trimmed mean of M values (TMM) method. We evaluated gene expression differences as a function of seasonal period and maternal body condition in a non-intercept model (∼0 + group). The model design was implemented with the *lmFit* and *eBayes* functions applied to contrasts of interest after controlling the variance with voom (R package limma v3.44.3) (Ritchie et al. 2015). These analysis steps were chosen following current best practices (Law et al. 2016, 2020).

#### Shared significant DEGs with previous diapause studies

We compared genes responding to time of season or body condition in our study and those associated with diapause in other species. We compared our genes of interest with those differentially expressed in diapausing queens in *B. terrestris* (supplement table S3D in Amsalem et al. (2015)), those related to diapause in larvae of the tropical oil-collecting bee *T. diversipes* (Supplement Table S1 and S2 in Santos et al. (2018)), as well as those related to early and late diapause induction in pupae of *M. rotundata* in November (supplement S2 in Yocum et al. (2018)). We identified orthologs as reciprocal best BLAST hits (evalue cutoff of 10×10^−5^), using amino acid sequences from Mrot v1.1 and amino acid sequences converted from nucleotide sequences with TransDecoder for *T. diversipies* or nucleotide sequences *B. terrestris*. We calculated representation factors and assessed significance of gene overlap using hypergeometric tests with the *phyper* function in R v. 4.2.1 (Hankin 2016).

#### Weighted Gene Co-expression Network Analysis

We conducted a weighted gene correlation network analysis (WGCNA) on our expression data to explore relationships between clusters of highly co-expressed genes (i.e., gene modules) and our experimental variables (Langfelder & Horvath 2008). To do so, we constructed scale-free gene co-expression networks using the R package WGCNA v1.70-3 (Langfelder & Horvath 2008) for mRNA expression in egg (*n*=22) and oocyte (*n*=22) samples independently. Using the default filtering parameters of the *goodSamplesGenes* function, we removed 852 genes from the expression data set due to excessive missing data or zero variance across samples (*n*=11,918 genes were retained for subsequent analyses). After filtering, we subset the data by sample type for tissue-specific analyses. We used hierarchical clustering (stats v3.6.1) to assess outliers. No outliers were identified among the oocyte samples, but two egg samples (202196 and 202198) were identified as outliers and removed (remaining: oocytes, *n* = 22; eggs, *n* = 20). Gene networks for oocytes and eggs were constructed at a soft power of 4, which appeared best suited for balancing scale independence (*R*^2^ = 0.624 and *R*^2^ = 0.529, respectively) and mean connectivity (303 and 458, respectively). In both cases, we computed a topological overlap matrix from the adjacency matrix using WGCNA default parameters.

Then we began module assignment. WGCNA gene modules are clusters of densely interconnected genes (Langfelder & Horvath 2008) defined here using an unsupervised hierarchical approach. Module membership for each gene is assigned using an eigengene-based connectivity approach; each gene is assigned to a single module by correlating the gene’s expression profile with the eigengene value of each module. Values close to 1 or -1 suggest high connectivity to a particular module (Langfelder & Horvath 2008). We initially assigned modules with a minimum module threshold of 30 genes and *deepSplit* =2. Modules with similar co-expression, i.e. those with a correlation *>* 75%, were subsequently merged using a cutHeight = 0.25. To quantify module-trait associations, we calculated the correlation of module eigengenes with cage (*n*=2), time of season (‘early’ vs ‘late’ season), and maternal condition (‘control’ vs ‘poor condition’) using Pearson correlation coefficient analysis (stats v3.6.1). Module-trait significance was assessed at a threshold of *α* = 0.05. Prior to making comparisons between oocytes and eggs, we matched and relabeled the egg network analysis module labels to the oocyte module labels by using oocyte labels as the reference (matchLabels(); reference = oocyte labels, source = egg labels; see Table S1 and S2).

We assessed module overlap between oocytes and eggs for those modules that were significantly associated with either season or body condition (henceforth termed ‘modules of interest’; *n*=4). One large egg module, *eggmod4*, had significant (*p <* 0.05) overlap with three of those oocyte modules of interest. We used bootstrapping methods to estimate the likelihood that this pattern of overlap could have occurred by chance. To do so, we randomly sampled (without replacement) gene ids from the list of all genes placed into module assignments equal to the number of genes in the respective modules of interest over 10,000 iterations. For each sample run, the proportion of genes sampled from *eggmod4* out of the total number of genes sampled was calculated. We then calculated the probability that the randomly sampled gene sets equal to the size of oocyte modules had a greater proportion of *eggmod4* genes than the observed proportion for each of modules of interest.

#### miRNA

miRNA expression was analyzed similar to the mRNA expression (R v4.0.2 R Core Team 2020). We used the matrix of counts to create a DGEList object with the EdgeR package v3.30.3 (Robinson et al. 2010). Visual inspection of unsupervised clustering plots (Fig. S1) revealed clear separation between oocyte and egg samples. We therefore proceeded to analyze these separately. For each dataset (oocyte and egg), we used the *filterByExpr* function to filter genes based on counts per million (CPM) and number of samples in each group. This kept 46 (61.3%) and 50 (66.7%) out of 75 miRNAs for oocytes and eggs, respectively. Libraries were then normalized with the TMM method. We evaluated gene expression differences as a function of time of season and maternal body condition in a non-intercept model (∼0 + group). The model design was implemented with the *lmFit* and *eBayes* functions applied to contrasts of interest after controlling the variance with voom (R package limma v3.44.3) (Ritchie et al. 2015). These analysis steps were chosen following current best practices (Law et al. 2016, 2020).

#### Gene Ontology Enrichment Analyses: mRNA and miRNA

Prior to enrichment analyses, we re-annotated the *M. rotundata* gene set v.1.1 (Kapheim et al. 2015) using InterProScan v5.56-89.0 (Jones et al. 2014) with Pfam (Mistry et al. 2020), SUPERFAMILY (Gough et al. 2001), and PANTHER (Thomas et al. 2022) analyses selected (-appl Pfam, SUPERFAMILY, PANTHER). Of the 12,770 genes, GO terms were assigned to 6,354 genes. Using those annotations, we performed Gene Ontology enrichment analyses for differentially expressed (Benjamini-Hochberg (BH)-adjusted *p <* 0.1) mRNA, predicted targets of differentially expressed miRNA (BH-adjusted *p <* 0.1), and the genes belonging to WGCNA modules of interest using TopGO v2.46.0 (Alexa & Rahnenfuhrer 2020) with default method (‘weight01’) and node size (nodesize=1) parameters.

## Results

### Differentially expressed mRNAs

We detected 8,276 genes in oocytes and 9,200 genes in eggs. Only 28.5% of genes detected in oocytes and 26.5% of genes detected in eggs have orthologs that were identified in a previous study of honey bee maternal RNA (Supplemental File 1) (Pires et al. 2016). Most of these had been previously classified as zygotic in origin (class II) in honey bees. Differences in maternal experience were evident in genes and miRNAs expressed in oocytes, but not in eggs (Table 1, Fig. 1). These effects were more apparent for differences in time of season than they were for maternal body condition (Fig. 1, Fig. S2). In oocytes, we detected 19 differentially expressed genes in response to poor body condition (8 up-regulated, 11 down-regulated; BH-adjusted *p <* 0.05). These included genes involved in embryogenesis, including orthologs of *branchless, knot*, and *juvenile hormone-inducible protein 26* (Supplemental File 1). To investigate additional genes that were less consistently differentially expressed, i.e., those missing the arbitrary *p <* 0.05 threshold, we relaxed *α* to BH-adjusted *p <* 0.1. Subsequently, the list grew to 35 differentially expressed genes (15 up-regulated, 20 down-regulated) (Table 1). These genes were enriched (*p <* 0.01) for enzyme regulator and proteasome-activating activity, as well as a transcription factor core complex (Table S3; Supplemental File 2). Seasonal effects in the oocyte were evident with 4 genes significantly up-regulated in the late season (BH-adjusted *p <* 0.05). These included genes involved in circadian rhythms and metabolic processes, including orthologs of *arylalkylamine N-acetyltransferase 1, knockdown*, and *chloride channel-b*. When we expanded *α* to BH-adjusted *p <* 0.1, we identified 413 genes up-regulated and 543 down-regulated in the late season (956 total). These genes were enriched (*p <* 0.01) for protein phosphorylation, dephosphorylation, and processing, regulation of transcription, and protein tyrosine phosphatase activity, as well as signal transduction and proton transmembrane transport (Table S4; Supplemental File 2). We did not detect any genes responding differently to body condition as a function of time of season, as would be indicated by the interaction term in our model. Only a single gene (Mrot07161) was identified as differentially expressed in eggs (Table 1). This was up-regulated in response to poor body condition. This result did not change when we adjusted the *α* to BH-adjusted *p <* 0.1. The *D. melanogaster* ortholog of Mrot07161 (*omega*) is involved in proteolysis, among other functions.

**Table 1.** Number of differentially expressed mRNAs and miRNAs at a threshold of a BH-adjusted *p* < 0.05 and (in parentheses) *p* < 0.1.

|  | mRNAs |  | miRNAs |  |
| --- | --- | --- | --- | --- |
|  | up | down | up | down |
| **Oocytes** |  |  |  |  |
| Poor condition | 8 (15) | 11 (20) | 0 (0) | 0 (0) |
| Late season | 4 (413) | 0 (543) | 2 (2) | 11 (17) |
| **Eggs** |  |  |  |  |
| Poor condition | 1 (1) | 0 (0) | 0 (0) | 0 (0) |
| Late season | 0 (0) | 0 (0) | 0 (0) | 0 (0) |

**Figure 1.**
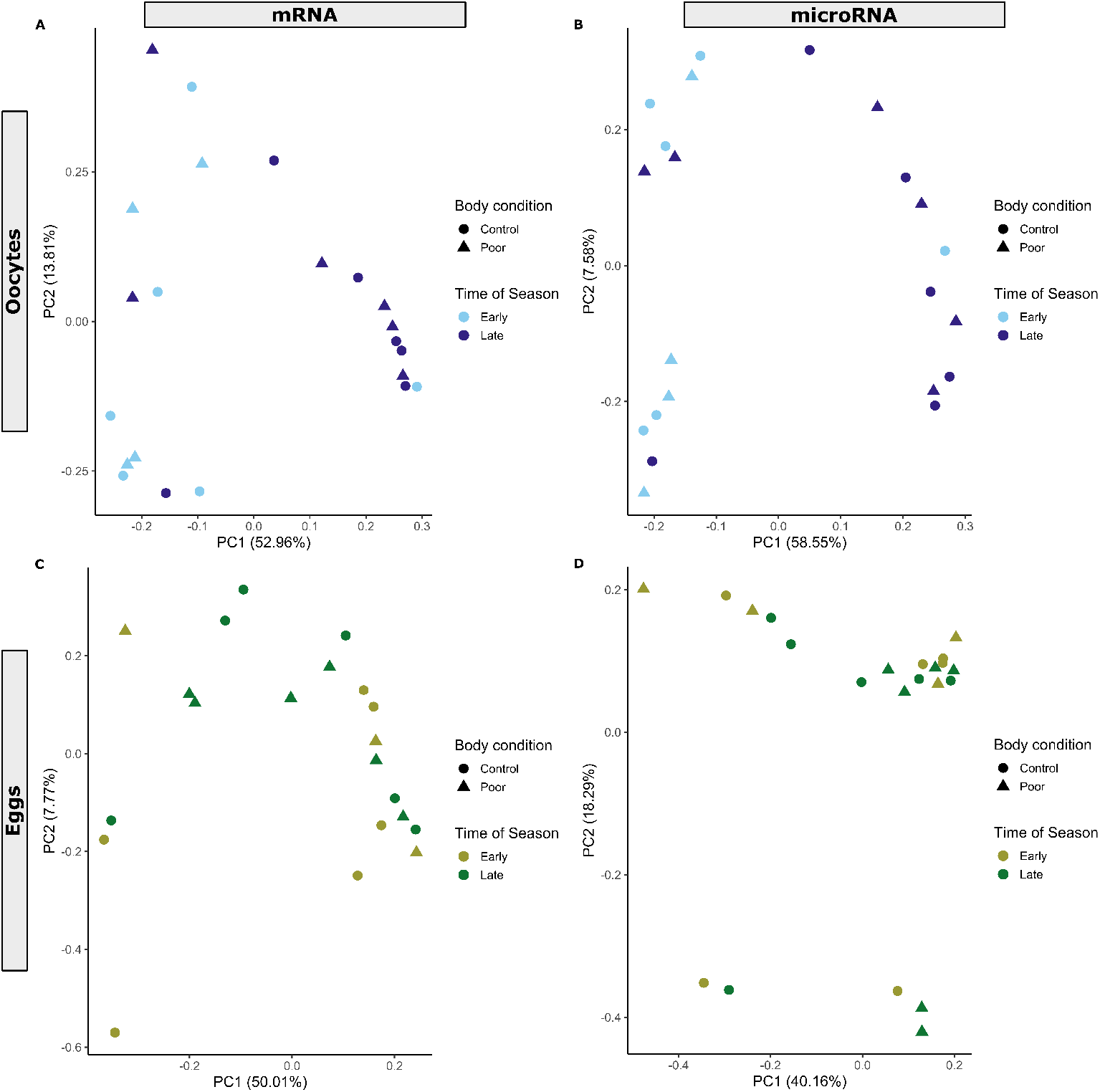
Principal component analysis showing mRNA and miRNA expression profiles for (A,B) oocyte (blue color scheme) and (C,D) egg (green color scheme) samples. Each data point represents an individual sample. Manipulations of maternal body condition included both control (circle) and poor condition (triangle) treatment groups.

Some of the maternal mRNAs that were differentially expressed in oocytes in response to maternal experience have been previously associated with the initiation or maintenance of diapause in insects. We compared genes differentially expressed between treatments in our oocyte tissue to those found differentially expressed in diapausing *B. terrestris* queens (Amsalem et al. 2015), diapausing *T. diversipes* larvae (Santos et al. 2018), and between fall-sampled *M. rotundata* prepupae that had entered diapause either in early and late summer (Yocum et al. 2018). The only significant overlap that we found was between our study and those related to diapausing *B. terrestris* queens (representation factor: 1.39; hypergeometric test: *p* = 0.07; Supplemental File 3).

### Differentially expressed miRNAs

More than half (39) of the 72 unique mature miRNA sequences identified in our oocyte and egg samples of *M. rotundata* were previously found in miRNA expression data from brains of six bee species (Kapheim et al. 2020). Many (34) of the 72 unique mature miRNA sequences identified in our oocyte and egg samples were previously found in miRNA expression data of *M. rotundata* brains (Fig. S3) (Kapheim et al. 2020). While only 72 unique mature miRNA sequences were identified, we characterized 75 distinct miRNAs for differential expression analysis. miRNAs with precursor sequences which did not overlap, but were paired with a mature miRNA that met all aforementioned precursor-mature pair filtering criteria were considered separate miRNAs for differential expression analysis, even if the mature sequence of one pair was identical to another. Of the 75 miRNAs expressed in our dataset, all except two had homologs in miRBase, and 80% (60) were detected in previous studies of maternal RNA or the MZT in insects (Supplemental File 4) (Marco 2015, Pires et al. 2016).

Differential expression analysis of miRNAs revealed a pattern similar to mRNAs (Fig. 1). Poor body condition did not yield any significant changes in miRNA expression in oocytes or eggs. We identified 13 miRNAs that responded significantly (BH-adjusted *p <* 0.05) to time of season in oocytes, with two up-regulated and 11 down-regulated in the late season, as compared to the early season (Table 1). While most of these 13 were involved in gene silencing, other functions of these miRNAs included regulation of the circadian rhythm, embryogenesis, and the cellular response to hypoxia (Supplemental File 4). Of these 13 miRNAs, 85% (11) were homologous to maternal miRNAs expressed in mature honey bee oocytes and degraded in early embryogenesis (Class I or III) (Pires et al. 2016), and 62% (8) were homologous to maternal miRNAs expressed in *D. melanogaster* oocytes (Marco 2015) (Supplemental File 4). We did not detect any miRNAs responding differently to body condition or time of season in the eggs (Table 1). Nor did we detect any miRNAs responding differently to body condition as a function of time of season in the oocytes or eggs, as would be indicated by the interaction term in our model.

We performed Gene Ontology enrichment of the set of predicted targets for the miRNAs differentially expressed across the season in the oocytes. This included a set of 164 unique genes, but there was no significant enrichment for any GO term (Supplemental File 5). There was no overlap between this set of predicted targets and the mRNA genes differentially expressed in oocytes in early and late season (BH-adjusted *p <* 0.1). However, more than half (56.3%) of the 375 unique predicted targets of miRNAs expressed in the eggs were genes expressed in oocytes, as would be expected if these miRNAs expressed in the early stages of embryogenesis function in clearing maternal transcripts (Marco 2015).

### Gene co-expression networks

The gene coexpression network for oocytes included 30 modules (Table S1, Supplemental File 6), none of which were significantly correlated with cage or maternal body condition (Fig. S4). However, we identified four modules (modules of interest) that were significantly associated with time of season (*oocyteMod4, r* = -0.57, p = 0.005; *oocyteMod29, r* = -0.51, p = 0.02; *oocyteMod16, r* = -0.46, p = 0.03; *oocyteMod20, r* = -0.45, p = 0.04; Figure S4). Genes belonging to these modules were enriched for various aspects of proton transmembrane transport, cell differentiation and cell fate determination, semaphorin receptor binding, vesicle-mediated transport, and tRNA activities, various types of substrate binding, and regulation of transcription (Table S5-S8, Supplemental File 7). The gene coexpression network for eggs included 27 modules, none of which were significantly correlated with cage, maternal body condition, or time of season (Table S2; Fig. S5; Supplemental File 8).

To investigate how the function of maternal RNAs shifts across the MZT, we evaluated the degree of overlap between gene coexpression modules in the oocytes and the eggs. We found that the structure of oocyte modules were not maintained in eggs (Fig. 2). However, one large egg module was more likely to include genes from the four oocyte modules of interest than expected by chance. Three oocyte modules (*oocyteMod16, oocyteMod4*, and *oocyteMod29*) had significant overlap with *eggMod4*, with 51.9% (*p* = 2.08*x*10^−20^), 45.1% (*p* = 0.00114), and 65.2% (*p* = 6.84*x*10^−10^) of their genes being assigned to this module in egg tissue, respectively (Table S9, Fig. S6, Supplemental File 9). Permutation tests suggest this degree of overlap was unlikely due to the large size of *eggMod4* alone (*p <* 0.0001 for *oocyteMod16, p <* 0.0016 for *oocyteMod4*, and *p <* 0.0001 for *oocyteMod29*).

**Figure 2.**
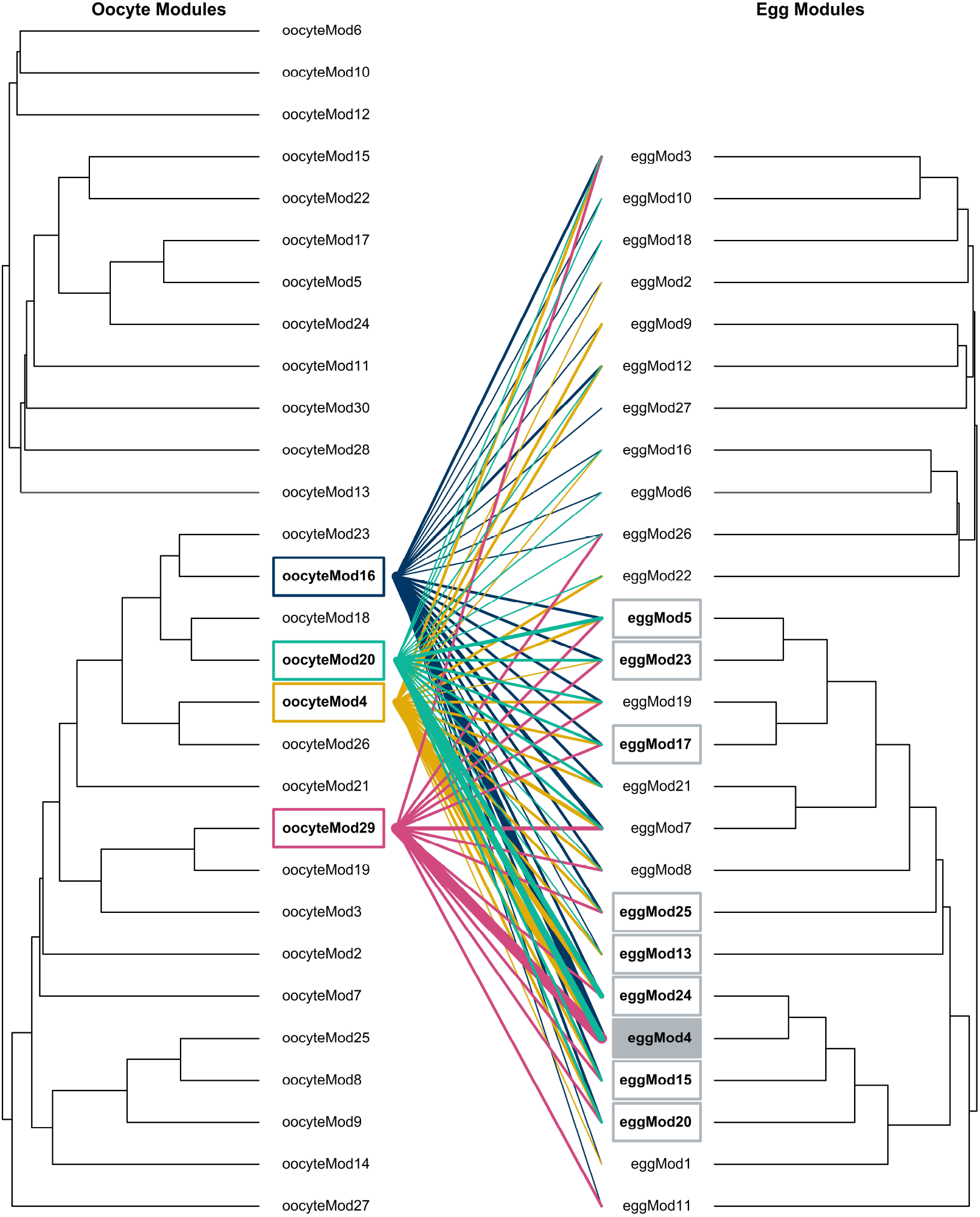
Dendrograms depicting gene module relationships for oocyte and egg tissues. Lines (moving left to right) indicate where genes from oocyte modules significantly correlated with photoperiod (*oocyteMod4, oocyteMod16, oocyteMod20, oocyteMod29*) are distributed throughout egg modules. The line color represents oocyte module membership, including gold, dark blue, turquoise, and magenta as *oocyteMod4, oocyteMod16, oocyteMod20, oocyteMod29*, respectively. Egg modules that shared significant overlap across these four oocyte modules are indicated with a gray box. The shaded box highlights (*eggMod4*), the egg module that shared a significant number of genes with three of these oocyte modules (*oocyteMod16, oocyteMod4*, and *oocyteMod29*). Line thickness scales to the percentage of genes from the oocyte modules that belong to each respective egg module. Line scale (from smallest to largest): <1%, 1-10%, 10-20%, 20-30%, 30-40%, 40-50%, 50-60%, and 60-70%.

## Discussion

Maternal effects are an important source of phenotypic diversity, but the mechanisms by which females interpret cues from the environment and translate them into molecular signals that influence offspring development are understudied. We found that alfalfa leafcutting bee females respond to cues from both the internal and external environment by adjusting the composition of RNA they deposit into maturing oocytes. These adjustments included both mRNAs and miRNAs previously known to be maternally-derived and playing a role in the earliest stages of development in insects. These effects were no longer detectable within 24 hours after egg-laying, which is consistent with current understanding of the maternal-to-zygotic transition. Our results shed light on the dynamic nature of how maternal experience can influence the developmental trajectory of her offspring.

Maternal mRNA provisioning is one of the earliest ways that a female can influence the phenotype of her offspring. These molecular signals are required for zygote genome activation in all animals (Berry 1982, Lee et al. 2014) and have been shown to affect survival after egg fertilization in mice (Christians et al. 2000), cell differentiation in nematodes (Mello et al. 1992), polarity in both fruit flies and nematodes (Evans et al. 1994), and cell fate determination in frogs (Xanthos et al. 2001). Given these large, organizational effects on offspring development, we hypothesized that adjustments to maternal RNA composition or abundance is an important mechanism underlying maternal effects. Indeed, research with mosquitoes and locusts suggests that female insects can influence offspring diapause outcome by adjusting mRNA provisioning of maturing oocytes in response to environmental cues (Poelchau et al. 2011, Hao et al. 2019). We found evidence that female alfalfa leafcutting bees also make dynamic adjustments to maternal RNAs, suggesting this could be a widespread mechanism by which females alter offspring development using cues from the environment. This is compelling, because *M. rotundata* diapause at a later stage of development (prepupae) than mosquitoes and locusts, which both diapause as pharate larvae inside the egg chorion. Our study reveals that development of later-diapausing species may also be influenced by maternal effects on the oocytes.

Given the role of maternal RNAs in processes critical to development (i.e., polarity and cell differentiation), we anticipated relatively few of these genes to be responsive to environmental cues. Instead, we anticipated most would have a narrow range of expression constrained by their role in canonical functions. This is consistent with our results, whereby fewer than 20 mRNAs and miRNAs showed highly significant differences in expression in response to female body condition or time of season. However, far more genes had less consistent or lower magnitude changes in expression, as indicated by the larger number of genes with significant differences in expression at a higher *α* (BH-adjusted *p <* 0.1; Table 1). These genes were enriched for functions that indicate high-level organizational effects on cellular differentiation, including processes related to transcription and translation. Thus, changes in the expression of even a small number of these genes could have potentially large effects on developmental trajectory. Notably, our assay only detected mRNAs active at the time of sample collection. Maternal RNA expression is dynamically regulated leading up to the MZT by the shortening and lengthening of polyA tails, a process that silences and activates translation, respectively (Winata & Korzh 2018). In this study, we used bead selection of polyA tails to generate RNA libraries for transcriptome sequencing. Hence, our results only capture the active portion of the maternal RNAs being passed to offspring. Additionally, we were unable to detect any adjustments that females made to the relative abundance of transcripts that were silenced until translation at later stages of the MZT. Thus, our results present a conservative picture of the dynamic changes possible for maternal RNAs.

The highly canonical role of maternal RNAs in development also suggests that the genes involved would be highly conserved across species. We found this to be the case for the miRNAs we detected in our samples, but less so for the mRNAs. Less than a third of the genes we detected in alfalfa leafcutting bee oocytes and eggs have orthologs that were identified in a previous study of honey bee maternal RNA (Pires et al. 2016). In contrast, the vast majority of the miRNAs we detected have homologs that were previously characterized as maternal in origin in honey bees or fruit flies (Marco 2015, Pires et al. 2016). One potential explanation for this is that there are fewer conserved protein-coding mRNAs than miRNAs in the *M. rotundata* genome, making it more likely to detect conserved miRNAs due to chance. However, the *M. rotundata* genome has roughly equal proportions of lineage-specific miRNAs and protein-coding genes (Kapheim et al. 2015, 2020). An alternative explanation is that the early developmental processes regulated by maternally-deposited miRNAs are more conserved than those regulated by maternally-deposited mRNAs. Across organisms, maternal miRNAs act, in part, to clear maternal mRNAs during the MZT by targeting maternal mRNA transcripts that were active during oocyte maturation, but are no longer needed during embryogenesis (Bashirullah et al. 1999, Winata & Korzh 2018). For example, in *D. melanogaster* the zygotically-expressed miR-309 cluster targets approximately 400 maternal mRNAs for degradation (Bushati et al. 2008). Consistent with this function, we found that more than half of the predicted targets of the miRNAs expressed in our 24-hr eggs were genes expressed in the oocytes. Interestingly, this suggests that the function of maternal miRNAs are conserved between alfalfa leafcutting bees and other insects (Marco 2015).

Conversely, we did not find evidence that maternal miRNAs and mRNAs share a coordinated response to environmental cues. There was no overlap between oocyte genes that were differentially expressed according to time of season and the predicted targets of oocyte miRNAs that were differentially expressed according to time of season. One potential explanation for this is a temporal lag in when these differences in expression would be detected. For instance, because it takes time for new target mRNAs to accumulate in the cytoplasm, a decrease in a mature miRNA might not correlate with an increase in target mRNAs. Thus, sampling mRNA profiles at additional time points would likely be necessary to observe the effects of environmentally-induced changes in miRNA expression on its target genes. However, our results are also consistent with previous work in *Drosophila* showing that only a small proportion of maternal transcripts are targeted by miRNAs during oocyte maturation (Nakahara et al. 2005). Further, there is some evidence that maternal transcripts have been selected to avoid targeting by maternal miRNAs, likely to preserve their function in oocyte maturation (Marco 2015).

Although our results offer compelling evidence that females adjust the mRNA and miRNA transcripts that they deposit into maturing oocytes in response to environmental cues, how this translates into offspring phenotype remains an open question. Answering this question is necessary to determine whether variation in maternal RNA composition is a mechanism underlying maternal effects. There is some evidence in various species of fish that variation in maternal RNAs alters the developmental trajectory of the embryos, resulting in different offspring phenotypes (Romney & Podrabsky 2017, Ahi et al. 2018, Torres-Paz et al. 2019). In mosquitoes, maternal adjustments to oocyte mRNA are concomitant with increased diapause incidence in offspring (Poelchau et al. 2011), and in locusts, knockdowns of maternal genes in the FOXO pathway have been causally linked to changes in offspring diapause (Hao et al. 2019). While direct evaluation of offspring phenotypes was outside the scope of our study, our results provide indirect evidence that variation in maternal RNAs in response to environmental cues could shape the developmental trajectory of alfalfa leafcutting bees. This evidence comes from what is already known about maternal effects in alfalfa leafcutting bees.

Whether leafcutting bee larvae enter diapause or not has long been recognized as being regulated by maternal effects (Tepedino & Parker 1986, Parker & Tepedino 1982, Bitner 1976, Kemp & Bosch 2001, Johansen & Eves 1973, Rank & Goerzen 1982), potentially triggered by changes in photoperiod (Pitts-Singer 2020, Wilson et al. 2021). In a recent study using an experimental design similar to that used in this study, we found that the proportion of offspring entering diapause was significantly higher among females nesting in the late season and significantly lower among females in poor body condition following experimental reduction of lipid stores (Johnson 2022). However, the mechanism regulating these effects appeared to vary. Females in poor body condition made smaller food provisions for their larvae, and size of food provision was a strong predictor of the diapause trajectory for individual larvae (Johnson 2022, Fischman et al. 2017). This could indicate that post-zygotic mechanisms of maternal effects (i.e., nutrient provisioning) acts in concert with pre-zygotic effects (i.e., maternal RNA provisioning) to influence offspring phenotype.

These differences in the size of larval food provisions were not evident between early and late season females, suggesting that the increased propensity for late season females to produce more diapausing offspring was more heavily influenced by pre-zygotic effects (i.e., maternal RNA provisioning) (Johnson 2022). Leafcutting bee larvae develop inside leaf-wrapped cell cups inside dark cavities, and are therefore unable to sense photoperiod directly. Moreover, changes in temperature have been ruled out as potential cues that regulate diapause (Pitts-Singer 2020, Wilson et al. 2021). While we cannot rule out other factors that correlate with time of season, such as changes in nutrient availability, these patterns are highly consistent with our findings—time of season leads to substantial significant differences in how females deposit mRNA and miRNA transcripts into maturing oocytes, but fewer significant differences associated with female body condition. Moreover, the genes differentially expressed in response to time of season had significant overlap with genes involved in diapause in another bee species and included genes and miRs with a known role in circadian rhythms and metabolism. Together, these results suggest that photoperiod-induced differences in maternally-provisioned RNA influence offspring propensity for diapause. Similar to this, environmentally-responsive maternal programming via mRNAs and miRNAs deposited in annual killifish oocytes determined whether young embryos develop directly or enter diapause (Romney & Podrabsky 2017). Overall, this suggests that diapause, as a developmental process, may be particularly sensitive to variation in maternal RNAs across species.

In contrast to oocytes, there was virtually no signal of variation in maternal environment observed in 24-hr eggs. This seems to suggest that whatever transcriptional adjustments females make to maturing oocytes do not persist beyond the MZT. This calls into question whether maternal RNAs are likely to have lasting effects on offspring phenotype. One possible explanation for this pattern is that the biochemical signatures of early embryogenesis dominated the transcriptomic profiles of 24-hr eggs, making any signal of maternal effects either difficult to detect or non-existent. For example, previous research in butterflies suggests that hormone pulses demarcating developmental transitions elicit a greater influence on gene expression profiles than environmental influences, such as temperature (Tian & Monteiro 2022). Our WGCNA analyses also showed signs of embryogenesis dominating transcriptional signatures in 24-hr eggs. While four modules of coexpressed genes correlated significantly with time of season in the oocytes, there were none in the eggs that correlated with either treatment. Further, when comparing patterns of co-regulated genes, major reorganization occurs between oocytes and eggs, such that no module of co-regulated genes remained entirely intact when progressing from oocyte to egg (Fig. 2). However, the fact that a significant proportion of genes belonging to the oocyte modules that responded to seasonal effects were coexpressed in the same egg module (*eggMod4*) could suggest that the maternal signal in oocytes is at least partially conserved in eggs. *eggMod4* was exceptionally large, with the genes from those four oocyte modules only comprising about one-tenth of those belonging to it. Yet, this is still more than was expected by chance (see permutation tests). Together, these results suggest that, while molecular signatures of maternal experience did not appear to be maintained past the MZT (i.e., in the egg), certain patterns of co-regulated genes may have persisted undetected against the overwhelming signal of transcriptional processes associated with embryogenesis.

However, it is also possible that the organizational effects of differential expression in maturing oocytes are no longer evident in transcriptomic data, but would be most readily observed in downstream processes. As such, effects of differentially expressed maternal RNAs in maturing oocytes might be more evident by examining metabolomic, proteomic, or endocrine profiles of eggs if expression changes have altered protein or metabolic production post-MZT. Future studies that incorporate additional types of biochemical data will be informative for understanding the effects of differences in maternal RNA expression on embryogenesis.

To understand how maternal experiences are translated into developmental changes in offspring requires investigating the molecular mechanisms underlying maternal effects. We found that both internal and external environmental cues elicit changes in mRNA and miRNA deposition by female bees into their maturing oocytes. This is consistent with our prior work showing that maternal photoperiod and body condition impacts offspring diapause induction in this species. Our current findings reveal how those environmental signals may facilitate facultative diapause in an economically important pollinator. Dynamic adjustment of maternal RNA in response to environmental cues has previously been documented in egg-diapausing insects, but it has heretofore been unclear whether similar mechanisms operate in species that diapause at a later life stage. Thus, our results also provide new insights about insect developmental biology, as well as mechanisms contributing to phenotypic variation in a changing world.

## Author Contributions

Conceptualization: K.M.K., T.L.P.S.; Data curation: M.A.H., F.K.H.; Formal analysis: M.A.H., F.K.H., T.J.D., K.M.K.; Funding acquisition: K.M.K., T.L.P.S.; Investigation: M.A.H., F.K.H., M.M.J.; Methodology: K.M.K., T.L.P.S.; Project administration: K.M.K.; Resources: K.M.K., T.L.P.S.; Supervision: K.M.K.; Visualization: M.A.H, K.M.K; Writing - original draft: M.A.H, F.K.H., T.J.D., K.M.K; Writing - Review & Editing: M.M.J, T.L.P.S.

## Acknowledgments

We are grateful to the farmers for cultivating and allowing us access to their alfalfa fields. We thank E. Klomps and N.K. Boyle for valuable discussion and assistance in the field. We thank R.A. Weinstock for their feedback on early manuscript drafts. We thank K. Hartfelder, S.M. Strobel, M.C. Womack, L.L. Surber, J.R. Phillips, R.A. Weinstock for helpful discussion of our results. Thanks to P.K. Santos for translated protein FASTA files of *T. diversipes*.This research was supported by the U.S. Department of Agriculture, Agricultural Research Service. Mention of trade names or commercial products in this publication is solely for the purpose of providing specific information and does not imply recommendation or endorsement by the U.S. Department of Agriculture.

## Supplemental Data

Supplemental Data files are attached.

## Data Availability Statement

Snakefiles (Sequence Alignment and Quantification) and R markdown (Statistical Analysis) files are available at Github: www.github.com/kapheimlab/mrotundata maternaleffects RNA. Sequences have been deposited at NCBI (PRJNA887543).

