## Supplement for "Maternal body condition and season influence RNA deposition in the oocytes of alfalfa leafcutting bees (*Megachile rotundata*)"

### Supplementary Material

#### Supplementary Data

Nine supplemental data files accompany the published manuscript. These files will be uploaded to FigShare for permanent storage upon acceptance.

#### Supplementary Tables

Table S1: Oocyte tissue: key for comparing renamed, manuscript-specific and WGCNA-generated module naming patterns. WGCNA assignments correspond to module ids used in the manuscript. Egg modules were matched to oocyte modules.

| Manuscript Name | Matched Name | Assigned Name |
|---|---|---|
| oocyteMod1 | - | grey |
| oocyteMod2 | - | saddlebrown |
| oocyteMod3 | - | floralwhite |
| oocyteMod4 | - | midnightblue |
| oocyteMod5 | - | black |
| oocyteMod6 | - | mediumpurple3 |
| oocyteMod7 | - | orangered4 |
| oocyteMod8 | - | orange |
| oocyteMod9 | - | lightyellow |
| oocyteMod10 | - | ivory |
| oocyteMod11 | - | white |
| oocyteMod12 | - | bisque4 |
| oocyteMod13 | - | skyblue3 |
| oocyteMod14 | - | darkolivegreen |
| oocyteMod15 | - | brown4 |
| oocyteMod16 | - | grey60 |
| oocyteMod17 | - | royalblue |
| oocyteMod18 | - | yellow |
| oocyteMod19 | - | darkred |
| oocyteMod20 | - | darkturquoise |
| oocyteMod21 | - | darkgrey |
| oocyteMod22 | - | blue |
| oocyteMod23 | - | lightsteelblue1 |
| oocyteMod24 | - | darkorange2 |
| oocyteMod25 | - | paleturquoise |
| oocyteMod26 | - | cyan |
| oocyteMod27 | - | darkmagenta |
| oocyteMod28 | - | lightcyan1 |
| oocyteMod29 | - | violet |
| oocyteMod30 | - | sienna3 |

Table S2: Egg tissue: key for comparing renamed, manuscript-specific and WGCNA-generated module naming patterns. Egg module ids were matched and relabeled to the oocyte module labels, as indicated by the *matched name* column.

| **Manuscript Name** | **Matched Name** | **Assigned Name** |
|---|---|---|
| eggMod1 | bisque4 | lightsteelblue1 |
| eggMod2 | purple | mediumpurple3 |
| eggMod3 | white | steelblue |
| eggMod4 | lightsteelblue1 | blue |
| eggMod5 | blue | yellowgreen |
| eggMod6 | brown4 | bisque4 |
| eggMod7 | lightyellow | brown |
| eggMod8 | darkred | lightgreen |
| eggMod9 | pink | darkslateblue |
| eggMod10 | sienna3 | sienna3 |
| eggMod11 | red | darkorange2 |
| eggMod12 | floralwhite | darkorange |
| eggMod13 | darkgrey | lightcyan1 |
| eggMod14 | grey | grey |
| eggMod15 | orange | darkgey |
| eggMod16 | saddlebrown | thistle1 |
| eggMod17 | midnightblue | darkmagenta |
| eggMod18 | paleturquoise | plum2 |
| eggMod19 | magenta | darkturquoise |
| eggMod20 | cyan | greenyellow |
| eggMod21 | black | cyan |
| eggMod22 | orangered4 | floralwhite |
| eggMod23 | royalblue | plum1 |
| eggMod24 | darkturquoise | green |
| eggMod25 | grey60 | midnightblue |
| eggMod26 | turquoise | brown4 |
| eggMod27 | skyblue3 | thistle2 |

Table S3: Enrichment results ($p < 0.05$) for genes that were significantly differentially expressed with maternal body condition in oocyte tissue. For full results, see Supplemental File 2.

| GO.ID | Terms | Annotated | Significant | Expected | $p$-value |
|---|---|---|---|---|---|
| *Biological Processes:* | | | | | |
| GO:0006506 | GPI anchor biosynthetic process | 21 | 0 | 0.06 | 0.011 |
| GO:0006388 | tRNA splicing, via endonucleolytic cleavage and ligation | 3 | 0 | 0.01 | 0.011 |
| GO:0006488 | dolichol-linked oligosaccharide biosynthetic process | 8 | 0 | 0.02 | 0.011 |
| GO:0006457 | protein folding | 30 | 0 | 0.09 | 0.014 |
| GO:0006412 | translation | 194 | 0 | 0.56 | 0.015 |
| GO:0071526 | semaphorin-plexin signaling pathway | 3 | 0 | 0.01 | 0.022 |
| GO:0003341 | cilium movement | 6 | 0 | 0.02 | 0.026 |
| GO:0006122 | mitochondrial electron transport, ubiquinol to cytochrome c | 5 | 0 | 0.01 | 0.032 |
| GO:0030488 | tRNA methylation | 3 | 0 | 0.01 | 0.032 |
| GO:0090263 | positive regulation of canonical Wnt signaling pathway | 2 | 0 | 0.01 | 0.034 |
| GO:0032049 | cardiolipin biosynthetic process | 2 | 0 | 0.01 | 0.035 |
| GO:0034220 | ion transmembrane transport | 70 | 0 | 0.2 | 0.036 |
| GO:0000079 | regulation of cyclin-dependent protein serine/threonine kinase activity | 3 | 0 | 0.01 | 0.039 |
| GO:0006508 | proteolysis | 210 | 1 | 0.61 | 0.042 |
| *Molecular Function:* | | | | | |
| GO:0036402 | proteasome-activating activity | 5 | 0 | 0.02 | 0.00014 |
| GO:0030234 | enzyme regulator activity | 141 | 1 | 0.51 | 0.0002 |
| GO:0016887 | ATP hydrolysis activity | 60 | 0 | 0.21 | 0.01022 |
| GO:0140662 | ATP-dependent protein folding chaperone | 19 | 0 | 0.07 | 0.01109 |
| GO:0004252 | serine-type endopeptidase activity | 41 | 0 | 0.15 | 0.01246 |
| GO:0015267 | channel activity | 78 | 0 | 0.28 | 0.01818 |
| GO:0003735 | structural constituent of ribosome | 120 | 0 | 0.43 | 0.01852 |
| GO:0017154 | semaphorin receptor activity | 3 | 0 | 0.01 | 0.02227 |
| GO:0016538 | cyclin-dependent protein serine/threonine kinase regulator activity | 12 | 0 | 0.04 | 0.02559 |
| GO:0019843 | rRNA binding | 14 | 0 | 0.05 | 0.0301 |
| GO:0051879 | Hsp90 protein binding | 5 | 0 | 0.02 | 0.03051 |
| GO:0004867 | serine-type endopeptidase inhibitor activity | 11 | 0 | 0.04 | 0.03167 |
| GO:0060072 | large conductance calcium-activated potassium channel activity | 3 | 0 | 0.01 | 0.03183 |
| GO:0004376 | glycolipid mannosyltransferase activity | 5 | 0 | 0.02 | 0.03477 |
| GO:0016772 | transferase activity, transferring phosphorus-containing groups | 322 | 1 | 1.15 | 0.03799 |
| GO:0016798 | hydrolase activity, acting on glycosyl bonds | 40 | 0 | 0.14 | 0.03931 |
| GO:0005230 | extracellular ligand-gated ion channel activity | 24 | 0 | 0.09 | 0.0427 |
| GO:0004298 | threonine-type endopeptidase activity | 2 | 0 | 0.01 | 0.04404 |
| GO:0000213 | tRNA-intron endonuclease activity | 2 | 0 | 0.01 | 0.04765 |
| GO:0051082 | unfolded protein binding | 17 | 0 | 0.06 | 0.04893 |
| GO:0016757 | glycosyltransferase activity | 84 | 1 | 0.3 | 0.04894 |
| *Cellular Components:* | | | | | |
| GO:0000439 | transcription factor TFIIH core complex | 4 | 0 | 0.01 | 0.0059 |
| GO:0005576 | extracellular region | 46 | 2 | 0.16 | 0.0203 |
| GO:0019773 | proteasome core complex, alpha-subunit complex | 7 | 1 | 0.02 | 0.0208 |
| GO:0005839 | proteasome core complex | 14 | 1 | 0.05 | 0.0239 |
| GO:0062023 | collagen-containing extracellular matrix | 2 | 0 | 0.01 | 0.0273 |
| GO:0046658 | anchored component of plasma membrane | 2 | 0 | 0.01 | 0.0273 |
| GO:0005887 | integral component of plasma membrane | 9 | 0 | 0.03 | 0.0321 |
| GO:0005750 | mitochondrial respiratory chain complex III | 5 | 0 | 0.02 | 0.0334 |
| GO:0005840 | ribosome | 121 | 0 | 0.42 | 0.0486 |

Table S4: Enrichment results ($p < 0.05$) for genes that were significantly differentially expressed with time of season in oocyte tissue. For full results, see Supplemental File 2.

| GO.ID | Terms | Annotated | Significant | Expected | *p*-value |
|---|---|---|---|---|---|
| *Biological Processes:* | | | | | |
| GO:1902600 | proton transmembrane transport | 26 | 6 | 3.03 | 0.000028 |
| GO:0006468 | protein phosphorylation | 199 | 27 | 23.21 | 0.00087 |
| GO:0006355 | regulation of transcription, DNA-templated | 226 | 36 | 26.36 | 0.00136 |
| GO:0051056 | regulation of small GTPase mediated signal transduction | 14 | 1 | 1.63 | 0.00607 |
| GO:0006470 | protein dephosphorylation | 33 | 8 | 3.85 | 0.00611 |
| GO:0016485 | protein processing | 13 | 3 | 1.52 | 0.00907 |
| GO:0046488 | phosphatidylinositol metabolic process | 34 | 4 | 3.97 | 0.02429 |
| GO:0017121 | plasma membrane phospholipid scrambling | 2 | 2 | 0.23 | 0.02636 |
| GO:0006821 | chloride transport | 4 | 1 | 0.47 | 0.02705 |
| GO:0070588 | calcium ion transmembrane transport | 9 | 3 | 1.05 | 0.03141 |
| GO:0006897 | endocytosis | 12 | 1 | 1.4 | 0.03442 |
| GO:0010506 | regulation of autophagy | 2 | 2 | 0.23 | 0.03738 |
| GO:0006874 | cellular calcium ion homeostasis | 7 | 3 | 0.82 | 0.03857 |
| GO:0042176 | regulation of protein catabolic process | 5 | 0 | 0.58 | 0.04374 |
| GO:0032469 | endoplasmic reticulum calcium ion homeostasis | 2 | 1 | 0.23 | 0.04806 |
| *Molecular Function:* | | | | | |
| GO:0004725 | protein tyrosine phosphatase activity | 17 | 6 | 2.12 | 0.00028 |
| GO:0046961 | proton-transporting ATPase activity, rotational mechanism | 11 | 6 | 1.37 | 0.00122 |
| GO:0005198 | structural molecule activity | 161 | 10 | 20.12 | 0.00639 |
| GO:0004672 | protein kinase activity | 199 | 26 | 24.86 | 0.0122 |
| GO:0008483 | transaminase activity | 12 | 1 | 1.5 | 0.01805 |
| GO:0046912 | acyltransferase, acyl groups converted into alkyl on transfer | 3 | 2 | 0.37 | 0.02237 |
| GO:0004190 | aspartic-type endopeptidase activity | 4 | 2 | 0.5 | 0.02319 |
| GO:0030276 | clathrin binding | 5 | 1 | 0.62 | 0.0235 |
| GO:0017128 | phospholipid scramblase activity | 2 | 2 | 0.25 | 0.02746 |
| GO:0016307 | phosphatidylinositol phosphate kinase activity | 3 | 2 | 0.37 | 0.02791 |
| GO:0005247 | voltage-gated chloride channel activity | 3 | 1 | 0.37 | 0.03203 |
| GO:0016810 | hydrolase activity, acting on carbon-nitrogen (but not peptide) bonds | 22 | 1 | 2.75 | 0.03279 |
| GO:0004523 | RNA-DNA hybrid ribonuclease activity | 3 | 1 | 0.37 | 0.0337 |
| GO:0009922 | fatty acid elongase activity | 9 | 2 | 1.12 | 0.03582 |
| GO:0004867 | serine-type endopeptidase inhibitor activity | 11 | 0 | 1.37 | 0.0371 |
| *Cellular Components:* | | | | | |
| GO:0005856 | cytoskeleton | 89 | 8 | 8.73 | 0.008 |
| GO:0000139 | Golgi membrane | 9 | 1 | 0.88 | 0.018 |
| GO:0016020 | membrane | 744 | 85 | 73.02 | 0.02 |
| GO:0005765 | lysosomal membrane | 5 | 1 | 0.49 | 0.026 |
| GO:0031932 | TORC2 complex | 3 | 1 | 0.29 | 0.035 |
| GO:0000502 | proteasome complex | 21 | 0 | 2.06 | 0.038 |
| GO:0000159 | protein phosphatase type 2A complex | 3 | 2 | 0.29 | 0.039 |
| GO:0008180 | COP9 signalosome | 7 | 1 | 0.69 | 0.043 |
| GO:0033177 | proton-transporting two-sector ATPase complex, proton-transporting domain | 13 | 3 | 1.28 | 0.043 |
| GO:0005783 | endoplasmic reticulum | 61 | 9 | 5.99 | 0.045 |
| GO:0005802 | trans-Golgi network | 4 | 1 | 0.39 | 0.048 |

Table S5: Enrichment results ($p < 0.05$) for genes from oocyte samples that were assigned to *OocyteMod29* via WGCNA analysis. *OocyteMod29* was significantly associated with time of season in oocyte tissue. For full results, see Supplemental File 8.

| GO.ID | Terms | Annotated | Significant | Expected | *p*-value |
|---|---|---|---|---|---|
| *Biological Processes:* | | | | | |
| GO:1902600 | proton transmembrane transport | 29 | 7 | 0.25 | $2.20\times10^{-9}$ |
| GO:0045596 | negative regulation of cell differentiation | 1 | 1 | 0.01 | 0.0085 |
| GO:0001709 | cell fate determination | 1 | 1 | 0.01 | 0.0085 |
| GO:0001678 | cellular glucose homeostasis | 2 | 1 | 0.02 | 0.017 |
| GO:1904668 | positive regulation of ubiquitin protein ligase activity | 3 | 1 | 0.03 | 0.0254 |
| GO:0035434 | copper ion transmembrane transport | 3 | 1 | 0.03 | 0.0254 |
| GO:0006879 | cellular iron ion homeostasis | 4 | 1 | 0.03 | 0.0337 |
| GO:0006826 | iron ion transport | 4 | 1 | 0.03 | 0.0337 |
| GO:0000122 | negative regulation of transcription by RNA polymerase II | 5 | 1 | 0.04 | 0.042 |
| *Molecular Function:* | | | | | |
| GO:0046961 | proton-transporting ATPase activity, rotational mechanism | 11 | 6 | 0.09 | $1.10\times10^{-9}$ |
| GO:0030215 | semaphorin receptor binding | 6 | 2 | 0.05 | 0.001 |
| GO:0015078 | proton transmembrane transporter activity | 34 | 8 | 0.28 | 0.012 |
| GO:0005536 | glucose binding | 2 | 1 | 0.02 | 0.017 |
| GO:0004396 | hexokinase activity | 2 | 1 | 0.02 | 0.017 |
| GO:0005375 | copper ion transmembrane transporter activity | 3 | 1 | 0.02 | 0.025 |
| GO:0097027 | ubiquitin-protein transferase activator activity | 3 | 1 | 0.02 | 0.025 |
| GO:0010997 | anaphase-promoting complex binding | 3 | 1 | 0.02 | 0.025 |
| GO:0004861 | cyclin-dependent protein serine/threonine kinase inhibitor activity | 4 | 1 | 0.03 | 0.033 |
| GO:0008199 | ferric iron binding | 5 | 1 | 0.04 | 0.041 |
| GO:0016298 | lipase activity | 41 | 2 | 0.34 | 0.045 |
| *Cellular Components:* | | | | | |
| GO:0033177 | proton-transporting two-sector ATPase complex, proton-transporting domain | 13 | 3 | 0.09 | 0.0015 |
| GO:0016471 | vacuolar proton-transporting V-type ATPase complex | 2 | 2 | 0.01 | 0.0068 |
| GO:0000221 | vacuolar proton-transporting V-type ATPase, V1 domain | 1 | 1 | 0.01 | 0.0073 |
| GO:0033178 | proton-transporting two-sector ATPase complex, catalytic domain | 6 | 2 | 0.04 | 0.0269 |
| GO:0033179 | proton-transporting V-type ATPase, V0 domain | 4 | 1 | 0.03 | 0.0288 |

Table S6: Enrichment results ($p < 0.05$) for genes from oocyte samples that were assigned to *OocyteMod20* via WGCNA analysis. *OocyteMod20* was significantly associated with time of season in oocyte tissue. For full results, see Supplemental File 8.

| **GO.ID** | **Terms** | **Annotated** | **Significant** | **Expected** | ***p*-value** |
|---|---|---|---|---|---|
| *Biological Processes:* | | | | | |
| GO:0016192 | vesicle-mediated transport | 131 | 24 | 9.96 | 0.0038 |
| GO:0006429 | leucyl-tRNA aminoacylation | 2 | 2 | 0.15 | 0.0058 |
| GO:0018216 | peptidyl-arginine methylation | 7 | 3 | 0.53 | 0.0121 |
| GO:0006829 | zinc ion transport | 3 | 2 | 0.23 | 0.0164 |
| GO:0009058 | biosynthetic process | 859 | 72 | 65.31 | 0.0247 |
| GO:0006888 | endoplasmic reticulum to Golgi vesicle-mediated transport | 16 | 4 | 1.22 | 0.0287 |
| GO:0006750 | glutathione biosynthetic process | 4 | 2 | 0.3 | 0.0312 |
| GO:0006379 | mRNA cleavage | 4 | 2 | 0.3 | 0.0312 |
| GO:0002098 | tRNA wobble uridine modification | 10 | 3 | 0.76 | 0.0349 |
| GO:0006886 | intracellular protein transport | 83 | 11 | 6.31 | 0.0399 |
| GO:0006779 | porphyrin-containing compound biosynthetic process | 6 | 3 | 0.46 | 0.0491 |
| GO:0042176 | regulation of protein catabolic process | 5 | 2 | 0.38 | 0.0494 |
| *Molecular Function:* | | | | | |
| GO:0004363 | glutathione synthase activity | 2 | 2 | 0.14 | 0.0052 |
| GO:0004823 | leucine-tRNA ligase activity | 2 | 2 | 0.14 | 0.0052 |
| GO:0016274 | protein-arginine N-methyltransferase activity | 7 | 3 | 0.51 | 0.0105 |
| GO:0032549 | ribonucleoside binding | 3 | 2 | 0.22 | 0.0148 |
| GO:0003735 | structural constituent of ribosome | 120 | 15 | 8.66 | 0.0248 |
| GO:0015299 | solute:proton antiporter activity | 4 | 2 | 0.29 | 0.0283 |
| GO:0017025 | TBP-class protein binding | 4 | 2 | 0.29 | 0.0283 |
| GO:0070122 | isopeptidase activity | 10 | 3 | 0.72 | 0.0305 |
| GO:0008083 | growth factor activity | 18 | 4 | 1.3 | 0.0363 |
| GO:0005544 | calcium-dependent phospholipid binding | 5 | 2 | 0.36 | 0.0449 |
| GO:0003887 | DNA-directed DNA polymerase activity | 5 | 2 | 0.36 | 0.0449 |
| *Cellular Components:* | | | | | |
| GO:0000502 | proteasome complex | 21 | 6 | 1.52 | 0.014 |
| GO:0072546 | EMC complex | 3 | 2 | 0.22 | 0.015 |
| GO:0030008 | TRAPP complex | 5 | 3 | 0.36 | 0.028 |
| GO:0005869 | dynactin complex | 4 | 2 | 0.29 | 0.028 |
| GO:0005763 | mitochondrial small ribosomal subunit | 4 | 2 | 0.29 | 0.028 |
| GO:0005765 | lysosomal membrane | 5 | 2 | 0.36 | 0.045 |

Table S7: Enrichment results ($p < 0.05$) for genes from oocyte samples that were assigned to *OocyteMod16* via WGCNA analysis. *OocyteMod16* was significantly associated with time of season in oocyte tissue. For full results, see Supplemental File 8.

| **GO.ID** | **Terms** | **Annotated** | **Significant** | **Expected** | ***p*-value** |
|---|---|---|---|---|---|
| *Biological Processes:* | | | | | |
| GO:0006813 | potassium ion transport | 36 | 8 | 1.8 | 0.0039 |
| GO:0045893 | positive regulation of transcription, DNA-templated | 16 | 4 | 0.8 | 0.0069 |
| GO:0007160 | cell-matrix adhesion | 3 | 2 | 0.15 | 0.0073 |
| GO:0007186 | G protein-coupled receptor signaling pathway | 114 | 12 | 5.72 | 0.0189 |
| GO:0071805 | potassium ion transmembrane transport | 22 | 4 | 1.1 | 0.022 |
| GO:0006801 | superoxide metabolic process | 5 | 2 | 0.25 | 0.0226 |
| GO:0019752 | carboxylic acid metabolic process | 113 | 7 | 5.67 | 0.0248 |
| GO:0006811 | ion transport | 198 | 22 | 9.93 | 0.0322 |
| GO:0055085 | transmembrane transport | 355 | 29 | 17.8 | 0.0483 |
| GO:0060271 | cilium assembly | 29 | 6 | 1.45 | 0.0485 |
| *Molecular Function:* | | | | | |
| GO:0005242 | inward rectifier potassium channel activity | 2 | 2 | 0.12 | 0.0036 |
| GO:0005506 | iron ion binding | 64 | 10 | 3.83 | 0.0043 |
| GO:0005515 | protein binding | 1507 | 117 | 90.17 | 0.0047 |
| GO:0005509 | calcium ion binding | 120 | 15 | 7.18 | 0.0049 |
| GO:0020037 | heme binding | 67 | 10 | 4.01 | 0.006 |
| GO:0070840 | dynein complex binding | 3 | 2 | 0.18 | 0.0103 |
| GO:0004497 | monooxygenase activity | 61 | 11 | 3.65 | 0.012 |
| GO:0046983 | protein dimerization activity | 108 | 12 | 6.46 | 0.013 |
| GO:0050661 | NADP binding | 9 | 3 | 0.54 | 0.0136 |
| GO:0005267 | potassium channel activity | 25 | 7 | 1.5 | 0.0157 |
| GO:0004842 | ubiquitin-protein transferase activity | 66 | 7 | 3.95 | 0.027 |
| GO:0004930 | G protein-coupled receptor activity | 98 | 11 | 5.86 | 0.0312 |
| GO:0005201 | extracellular matrix structural constituent | 5 | 2 | 0.3 | 0.0316 |
| GO:0016705 | oxidoreductase activity, acting on paired donors, with incorporation or reduction of molecular oxygen | 72 | 10 | 4.31 | 0.0377 |
| *Cellular Components:* | | | | | |
| GO:0016021 | integral component of membrane | 519 | 44 | 29.21 | 0.0021 |
| GO:0005861 | troponin complex | 2 | 2 | 0.11 | 0.0031 |
| GO:0005581 | collagen trimer | 2 | 2 | 0.11 | 0.0031 |
| GO:0016020 | membrane | 1008 | 75 | 56.73 | 0.0122 |
| GO:0034464 | BBSome | 6 | 2 | 0.34 | 0.0406 |
| GO:0005576 | extracellular region | 73 | 8 | 4.11 | 0.0443 |

Table S8: Enrichment results ($p < 0.05$) for genes from oocyte samples that were assigned to *OocyteMod4* via WGCNA analysis. *OocyteMod4* was significantly associated with time of season in oocyte tissue. For full results, see Supplemental File 8.

| GO.ID | Terms | Annotated | Significant | Expected | $p$-value |
|---|---|---|---|---|---|
| *Biological Processes:* | | | | | |
| GO:0071929 | alpha-tubulin acetylation | 1 | 1 | 0.01 | 0.014 |
| GO:0043063 | intercellular bridge organization | 1 | 1 | 0.01 | 0.014 |
| GO:0002218 | activation of innate immune response | 1 | 1 | 0.01 | 0.014 |
| GO:0007300 | ovarian nurse cell to oocyte transport | 1 | 1 | 0.01 | 0.014 |
| GO:0032481 | positive regulation of type I interferon production | 1 | 1 | 0.01 | 0.014 |
| GO:0007140 | male meiotic nuclear division | 1 | 1 | 0.01 | 0.014 |
| GO:0006814 | sodium ion transport | 14 | 2 | 0.19 | 0.015 |
| GO:0007608 | sensory perception of smell | 43 | 3 | 0.59 | 0.021 |
| GO:0008608 | attachment of spindle microtubules to kinetochore | 2 | 1 | 0.03 | 0.027 |
| GO:0035434 | copper ion transmembrane transport | 3 | 1 | 0.04 | 0.041 |
| GO:1905515 | non-motile cilium assembly | 3 | 1 | 0.04 | 0.041 |
| GO:0006098 | pentose-phosphate shunt | 3 | 1 | 0.04 | 0.041 |
| *Molecular Function:* | | | | | |
| GO:0046961 | proton-transporting ATPase activity, rotational mechanism | 11 | 2 | 0.16 | 0.011 |
| GO:0009922 | fatty acid elongase activity | 12 | 2 | 0.18 | 0.013 |
| GO:0004801 | transaldolase activity | 1 | 1 | 0.01 | 0.015 |
| GO:0008113 | peptide-methionine (S)-S-oxide reductase activity | 1 | 1 | 0.01 | 0.015 |
| GO:0019799 | tubulin N-acetyltransferase activity | 1 | 1 | 0.01 | 0.015 |
| GO:0080019 | fatty-acyl-CoA reductase (alcohol-forming) activity | 13 | 2 | 0.19 | 0.015 |
| GO:0051015 | actin filament binding | 16 | 2 | 0.24 | 0.023 |
| GO:0004984 | olfactory receptor activity | 43 | 3 | 0.64 | 0.025 |
| GO:0140575 | transmembrane monodehydroascorbate reductase activity | 2 | 1 | 0.03 | 0.029 |
| GO:0004471 | malate dehydrogenase (decarboxylating) (NAD+) activity | 2 | 1 | 0.03 | 0.029 |
| GO:0004731 | purine-nucleoside phosphorylase activity | 2 | 1 | 0.03 | 0.029 |
| GO:0005549 | odorant binding | 50 | 3 | 0.74 | 0.037 |
| GO:0005375 | copper ion transmembrane transporter activity | 3 | 1 | 0.04 | 0.044 |
| GO:0003727 | single-stranded RNA binding | 3 | 1 | 0.04 | 0.044 |
| *Cellular Components:* | | | | | |
| GO:0033180 | proton-transporting V-type ATPase, V1 domain | 2 | 1 | 0.03 | 0.027 |
| GO:0005890 | sodium:potassium-exchanging ATPase complex | 3 | 1 | 0.04 | 0.04 |

Table S9: WGCNA modules for oocyte tissue that are significantly associated with time of season, and the corresponding modules for egg tissue with a significant number of overlapping genes. The number of genes in each oocyte module and their correlation with time of season (column 2), as well as the number of genes in each egg modules and their relationship with time of season (column 4) are provided. For oocyte module-egg module pairs, the amount of genes within each oocyte modules that overlap with each egg module are reported as percentages (i.e., of overlapping genes/ number of genes in each oocyte module). $p$-values and representation factors (RF) are given for each module-module pair. Asterisk indicate $p$-values of correlation with time of season: $*p < 0.05$, $**p < 0.01$, $**p < 0.001$

| | # genes ($r$ with season) | Egg Module | # genes ($r$ with season) | % Overlap (with Egg Module) | Overlap $p$-value | RF |
|---|---|---|---|---|---|---|
| OocyteMod16 | 588 (-0.46*) | eggMod25 | 230 (0.18) | 4.1 | 0.00043 | 2.115 |
| | | eggMod4 | 4031 (-0.28) | 51.9 | $2.08\text{x}10^{-20}$ | 1.534 |
| | | eggMod17 | 330 (-0.084) | 4.3 | 0.02191 | 1.536 |
| | | eggMod20 | 289 (0.079) | 3.6 | 0.049 | 1.473 |
| OocyteMod4 | 175 (-0.57**) | eggMod13 | 78 (0.2) | 2.3 | 0.02765 | 3.492 |
| | | eggMod4 | 4031 (-0.28) | 45.1 | 0.00114 | 1.335 |
| | | eggMod17 | 330 (-0.084) | 5.7 | 0.0238 | 2.064 |
| | | eggMod15 | 192 (-0.29) | 6.3 | 0.00012 | 3.902 |
| OocyteMod29 | 92 (-0.51*) | eggMod4 | 4031 (-0.28) | 65.2 | $6.84\text{x}10^{-10}$ | 1.928 |
| OocyteMod20 | 716 (-0.45*) | eggMod24 | 865 (-0.089) | 25 | $1.45\text{x}10^{-53}$ | 3.445 |
| | | eggMod5 | 645 (0.034) | 12 | $1.10\text{x}10^{-12}$ | 2.219 |
| | | eggMod23 | 287 (-0.18) | 4.2 | 0.00216 | 1.74 |

### Supplementary Figures

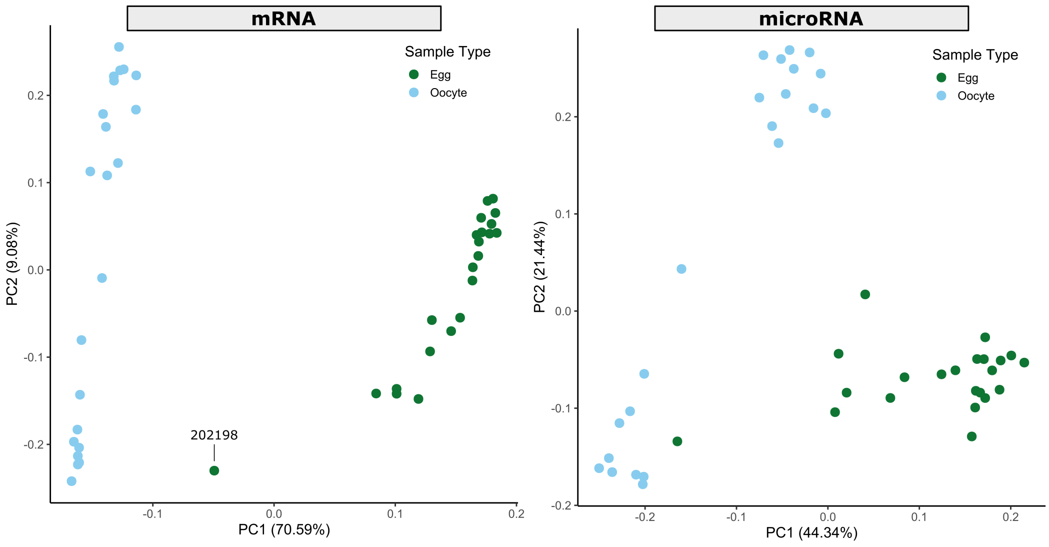


Figure 1: Principal component analysis (PCA) plots for mRNA and microRNA data showing variance between individual egg (green) and oocyte (blue) tissue samples. Egg samples were 24 h post-oviposition; oocytes were in stage 4 of maturation. Sample 202198 was identified as an outlier and removed from subsequent WGCNA analyses.

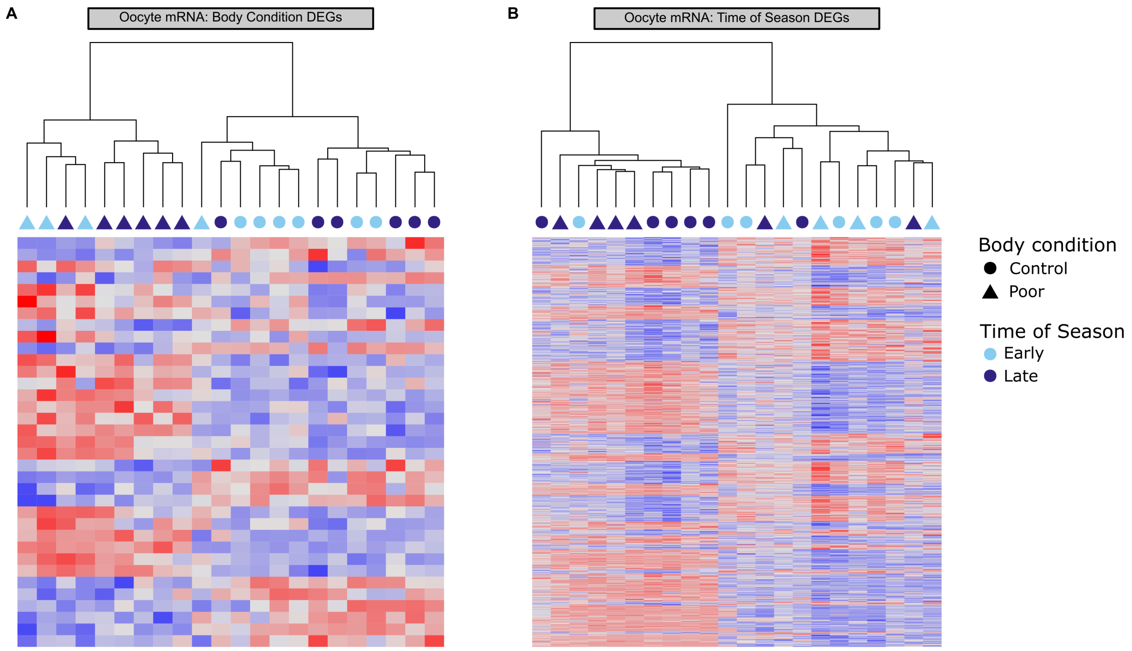


Figure 2: Significantly differentially expressed genes in oocyte tissue associated with (A) body condition and (B) photoperiod. Samples (x-axis) were either control (circle) or treatment (triangle) bees collected during the early (light blue) and late (dark blue) season. Expression key: up-regulated (red) and down-regulated (blue).

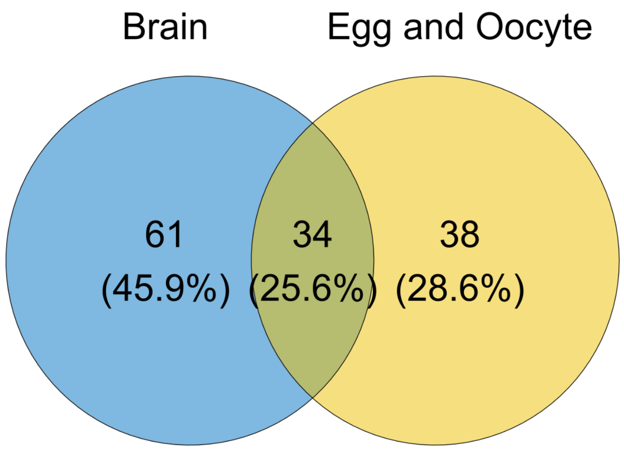


Figure 3: Venn Diagram depicting the overlap between *Megachile rotundata* brain miRNAs (Kapheim et al. 2020) and the egg/oocyte miRNAs identified in the current study.

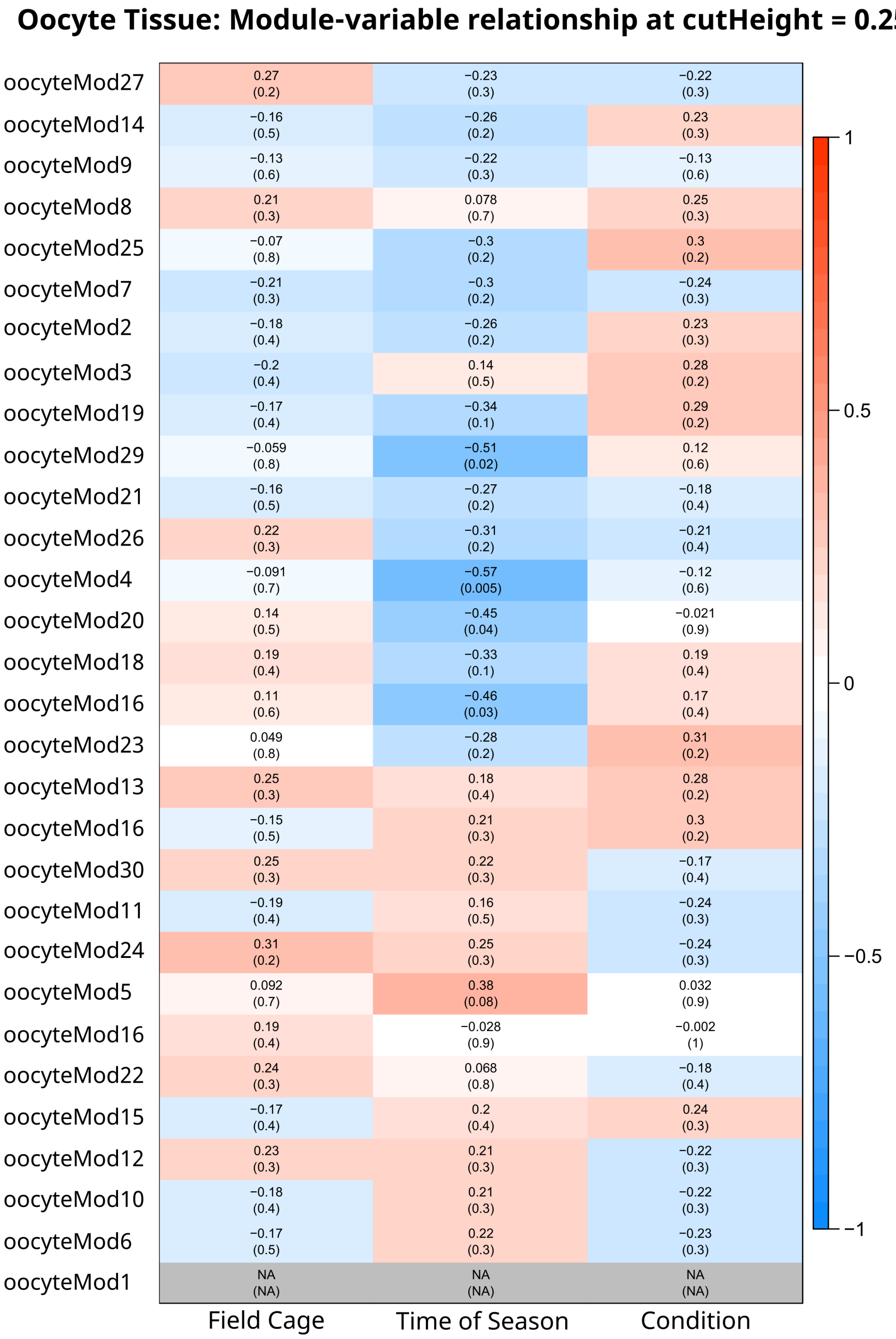


Figure 4: Correlation matrix between WGCNA gene modules (cut height = 0.25) and experimental variables for oocyte tissue. Correlation coefficients for each module-variable relationship, and their significance, are shown as r (p-value) and indicated using the color scale (positive relationship=red; negative relationship=blue; no relationship=white).

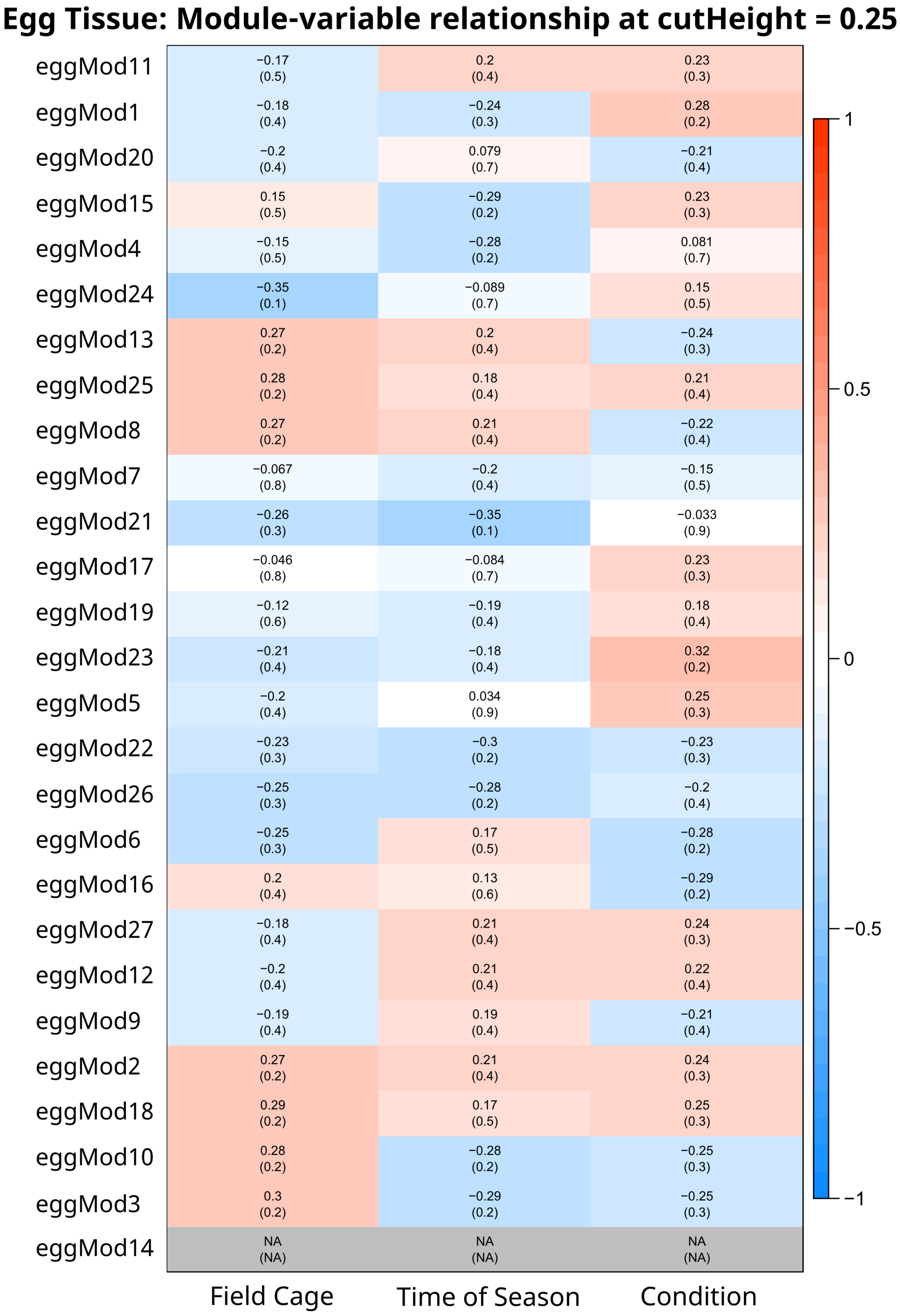


Figure 5: Correlation matrix between WGCNA gene modules (cut height = 0.25) and experimental variables for egg tissue. Correlation coefficients for each module-variable relationship, and their significance, are shown as r (p-value) and indicated using the color scale (positive relationship=red; negative relationship=blue; no relationship=white).

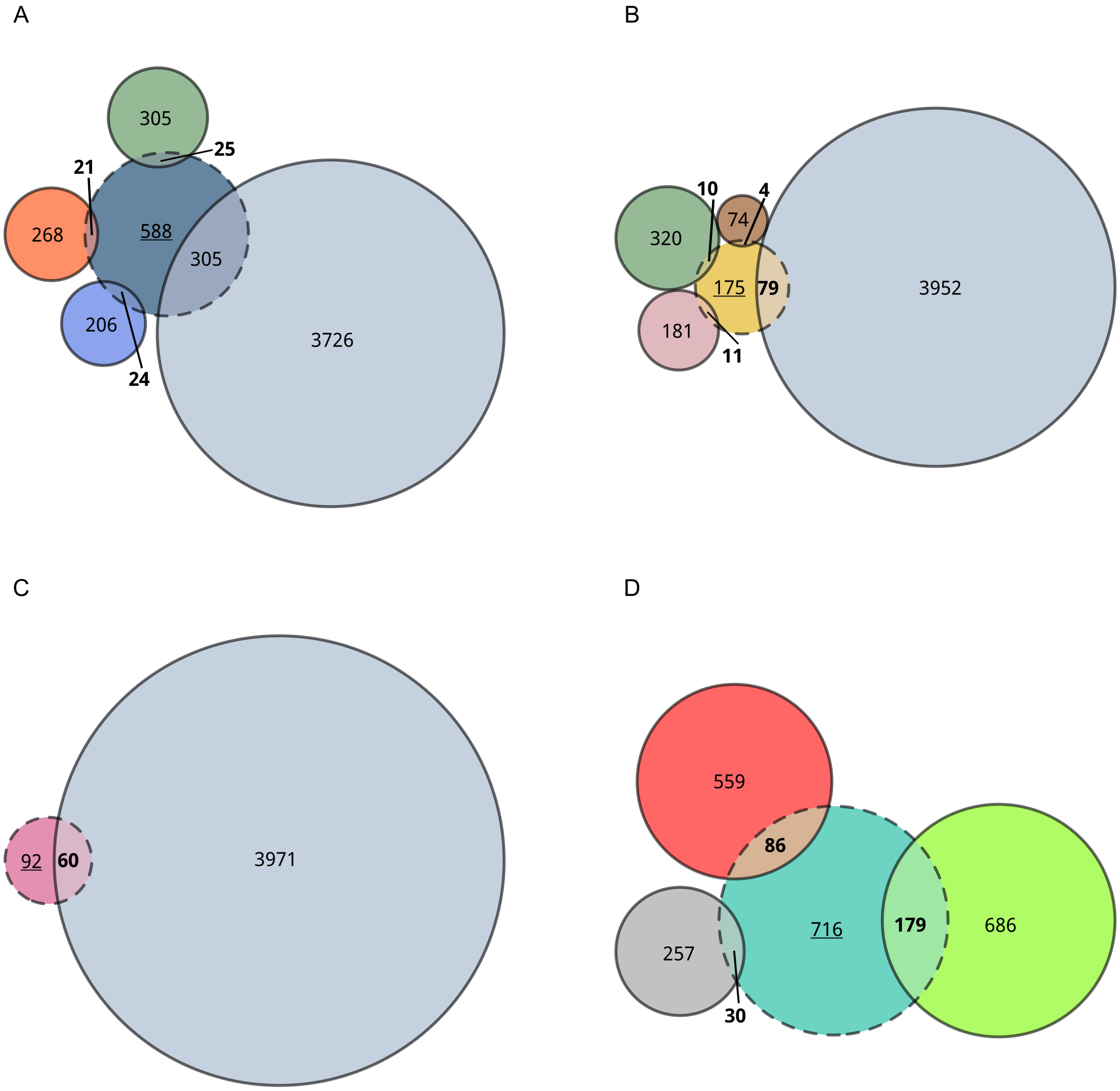


Figure 6: Oocyte modules of interest (i.e., correlated with time of season) and egg modules with which they have significant overlap. Oocyte modules, including (A) *oocyteMod16*, (B) *oocyteMod4*, (C) *oocyteMod29*, and (D) *oocyteMod20* are indicated using the dashed lines. The actual gene counts per oocyte module are underlined. Egg modules with significant overlap are indicated using solid lines and their gene counts, minus the overlap number, are present in their respective circle. The number of genes with overlap are bold and indicated at the overlapping space.
